# Specialised super-enhancer networks in stem cells and neurons

**DOI:** 10.1101/2025.08.13.670083

**Authors:** Izabela Harabula, Liam Speakman, Francesco Musella, Luca Forillo, Luna Zea-Redondo, Alexander Kukalev, Robert A Beagrie, Kelly J. Morris, Lucas Fernandes, Ibai Irastorza-Azcarate, Ana M. Fernandes, Silvia Carvalho, Dominik Szabó, Carmelo Ferrai, Mario Nicodemi, Lonnie Welch, Ana Pombo

**Author notes:** Department of Microbiology, Immunology, and Molecular Genetics, University of California Los Angeles, Los Angeles, California, USA. Institute of Quantitative and Computational Biosciences, University of California Los Angeles, Los Angeles, California, USA. Centre for Human Genetics, Roosevelt Drive Oxford, OX3 7BN, UK. Berlin Institute of Health at Charité-Universitätsmedizin Berlin, Center for Regenerative Therapies, Berlin, Germany (Silvia Carvalho). Max Planck Institute for Molecular Genetics, Berlin, Germany (Silvia Carvalho). Institute of Pathology, University Medical Center Göttingen, Robert-Koch-Straße 40, 37075 Göttingen, Germany. joint first authors (Izabela Harabula, Liam Speakman).

## Abstract

Super-enhancers (SEs) are clusters of enhancers with high transcriptional activity that play essential roles in defining cell identity through regulation of nearby genes. SEs preferentially form multiway chromatin interactions with other SEs and highly transcribed regions in embryonic stem cells. However, the properties of the interacting SEs and their specific contributions to complex regulatory interactions in differentiated cell types remain poorly understood. Here, we compare the structural and functional properties of SEs between embryonic stem cells (ESCs) and dopaminergic neurons (DNs) by combining Genome Architecture Mapping (GAM), chromatin accessibility, histone modification, and transcriptome data. Most SEs are cell-type specific and establish extensive pairwise and multiway chromatin interactions with other SEs and genes with cell-type specific expression. SE interactions span megabase genomic distances and frequently connect distant topologically associating domains. By applying network centrality analyses, we detected SEs with different hierarchical importance. Highest network centrality SEs contain binding motifs for cell-type specific transcription factors, and are candidate regulatory hubs. The functional heterogeneity of SEs is also highlighted by their organisation into modular sub-networks that differ in structure and spatial scale between ESCs and DNs, with more specific and strongly connected SE modules in post-mitotic neurons. Our results uncover both the high complexity and specificity of SE-based 3D regulatory networks and provide a resource for prioritizing SEs with potential roles in transcriptional regulation and disease.

## Introduction

Super-enhancers (SEs) were originally identified in mouse ESCs as clusters of enhancers with unusually high activity, which are densely occupied by cell-type specific master transcription factors (TFs) and the mediator complex (MED; Whyte et al. 2013). SEs often coincide with locus control regions (Gurumurthy et al. 2019), but are formally detected from genomic annotations based on ChIP-Seq enrichments for H3K27ac, H3K4me1, TFs, MED and/or other coactivators, using bioinformatics pipelines such as ROSE (Ranking of Super-Enhancers; (Lovén et al. 2013; Whyte et al. 2013). SEs are important for regulation of lineage-specific genes. For example, in ESCs, *Sox2* has essential roles in maintaining pluripotency and 90% of its expression is driven by a downstream SE (Li et al. 2014; Zhou et al. 2014). However, the function of SEs beyond assemblies of conventional enhancers with different regulatory properties remains controversial (Hay et al. 2016; Xie et al. 2017). Understanding SEs properties and their modes of action is of particular interest in the context of diseases such as cancer, as SEs can be used to predict metastasis progression and appear to be promising therapeutic targets (Lovén et al. 2013; Mack et al. 2018).

Several mechanisms have been proposed to explain how SEs regulate gene expression. Imaging-based technologies favour the condensate model, which proposes that SEs act as nucleation points for TFs (Sabari et al. 2018; Tang et al. 2022). For instance, in mouse ESCs, SEs physically associate with condensates of BRD4 (Bromodomain-contacting protein 4) and MED complex which can compartmentalize and concentrate the transcription machinery (Sabari et al. 2018). Technologies based on chromosome conformation capture (3C) detect local interactions between SEs and nearby promoters, suggesting that SEs regulate gene expression locally (Tang et al. 2022). However, complex multiway interactions across tens of megabases between SEs and highly transcribed genomic regions were identified, also in mouse ESCs, using Genome Architecture Mapping (GAM; (Beagrie et al. 2017; Beagrie et al. 2023), and validated by FISH (Beagrie et al. 2017) and by SPRITE (Quinodoz et al. 2018; Kempfer and Pombo 2020).

Chromatin interactions between enhancers and promoters are being extensively studied for their direct role in gene regulation in the context of development, homeostasis and disease. Enhancer-promoter interactions are currently thought to be predominantly confined within topologically associating domains (TADs), which are approx. 1 Mb genomic domains characterized by preferred local interactions delimited by boundary regions (Okhovat et al. 2023; Rajderkar et al. 2023). Inter-TAD regulatory chromatin contacts have been reported, for example at the *Dbx2* locus (Beccari et al. 2021), and may include mechanisms that depend on contacts between TAD boundaries (Hung et al. 2024). Megabase-long chromatin interactions between TADs have been detected genome-wide by Hi-C in mouse ESCs, NPCs and *in vitro* dopaminergic neurons (DNs) (Fraser et al. 2015) and by GAM in ESCs (Beagrie et al. 2017; Beagrie et al. 2023), in association with higher expression levels of the interacting regions. Megabase-range chromatin interactions between expressed genes have also been identified in neurons from adult mouse brains (Winick-Ng et al. 2021). Nonetheless, it remains unclear whether complex chromatin interactions involving many genomic partners become more prominent in post-mitotic cells, such as neurons, and which genomic regions and regulatory mechanisms drive the formation of complex hubs of SEs and highly expressed genes.

Here, we hypothesize that cell-type specific SEs form local and distal pairwise and three-way chromatin interactions with specific genes to regulate cell-type specific transcriptional programs (**Figure 1A**). To uncover the most prominent interacting partners of SEs, their specificity and their potential regulatory functions, we collected GAM from mouse ESCs and *in vitro* differentiated DNs, and integrated features of 3D genome organization with local chromatin accessibility, histone modifications and gene expression collected from the same cell types (**Figure 1B**). A summary of the datasets used in this study can be found in **Supplemental Table S1**. We show that most ESC and DN SEs were found to be cell-type specific. In both cell types, the most prominent chromatin interactions between SEs and differentially expressed genes (DEGs) reflected their cell-type specificity and specialization, and connected TADs separated by tens of megabases. We applied network centrality measures to explore the ESC– and DN-specific complex regulatory networks of SEs, rank SEs according to their network centrality, and identify the TF binding motifs most enriched in complex SE contacts. Our work suggests that SEs have specific functional and structural roles within the networks of chromatin interactions, and prioritizes SEs for further research according to their hierarchical contributions to the cell-type specific networks of chromatin contacts.

**Figure 1.**
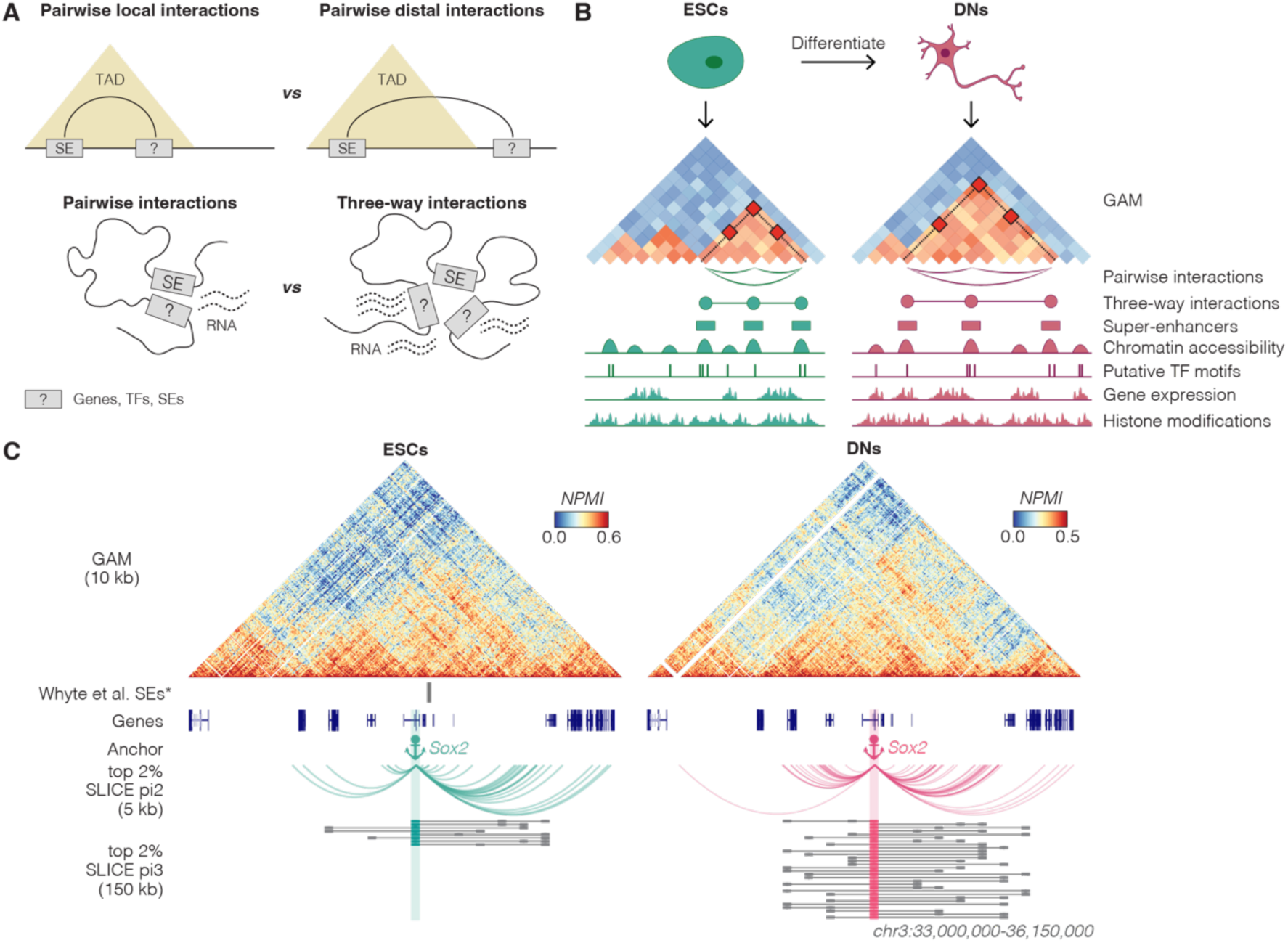
Summary of the key principles of the study. (**A**) Schematic representation of the main hypothesis and study aims. (**B**) Schematic representation of the study design, showing the cell types used, the datasets collected and the features derived. (**C**) Genome Architecture Mapping (GAM) matrices at 10 kb resolution around the *Sox2* gene locus. Arc plots show the top 2% significant pairwise SLICE *Pi2* interactions of the genomic window that overlaps *Sox2* at 5 kb resolution. Lollipop plots show the top 2% significant three-way SLICE *Pi3* interactions of the genomic window that overlaps *Sox2* at 150 kb resolution. *Represented Whyte et al. (2013) SEs exclude SEs that overlap promoters.

## Results

### Mapping long-range chromatin interactions with GAM in ESCs and DNs

Previous studies show an increased complexity in chromatin contacts across longer genomic distances between ESCs and terminally differentiated neurons (Fraser et al. 2015; Bonev et al. 2017; Winick-Ng et al. 2021). To find novel regulatory mechanisms that link chromatin topology to cellular identity and functions in specialized cells, we differentiated mouse ESCs into mature DNs (**Supplemental Figure S1A,B**). The differentiation system chosen yields a highly homogeneous DN population with the molecular and electrophysiological properties of mature DNs from the midbrain ventral tegmental area (VTA; day 30 neurons; Ferrai et al. 2017), important for reward, motivation, learning and addiction. ESCs are pluripotent and highly dividing cells, in contrast to DNs which are post-mitotic cells and have highly specialized roles that are sustained over the healthy lifespan of individuals.

To study the structural organization of SEs in terminally differentiated DNs, we produced GAM data from *in vitro*differentiated DNs (**Supplemental Figure S1C; GSE304996**), using multiplex immunoGAM (Beagrie et al. 2017; Winick-Ng et al. 2021). DNs were immuno-selected based on the detection of tyrosine hydroxylase (TH), a marker for mature DNs (Winick-Ng et al. 2021). We also re-processed a previously published GAM dataset from ESCs (Beagrie et al. 2023) into an equivalent 3NP GAM dataset. Quality control of both GAM datasets was performed in the individual biological replicates, before merging them, resulting in GAM datasets corresponding to 1,335 ESCs and 1,446 DNs (**Supplemental Figure S1D,E; Supplemental Tables S2-S4**). The merged datasets have high sampling quality in both ESCs and DNs, and enable contact mapping of pairwise interactions at 5 kb resolution, and of three-way interactions at 150 kb, the highest resolution achieved by GAM datasets to date.

### Identifying the most prominent two– and three-way contacts in ESCs and DNs

GAM is a technology that in combination with the mathematical model SLICE can detect significantly enriched chromatin interactions across all genomic distances (Beagrie et al. 2017; Beagrie et al. 2023). Here, we applied SLICE to detect pairwise and three-way interactions. SLICE infers the probability of pair-wise interactions (*Pi2*) between different genomic loci above their proximity expected from linear genomic distance and nuclear volume, by estimating the proportion interacting and non-interacting co-segregation frequencies that best explain the observed co-segregation of the respective loci (Beagrie et al. 2017; Beagrie et al. 2023). SLICE detected 318×10^6^ and 324×10^6^ significant pairwise intra-chromosomal interactions at 5 kb resolution in ESCs and DNs, respectively (**GSE304996**). We find that many genomic regions form hundreds of interactions in both ESCs and DNs (**Supplemental Figure S1F**).

To quantify three-way interactions that occur simultaneously in the same cell, SLICE considers all possible combinations of three 150 kb windows in each chromosome, and compares their experimental co-occurrence with their expected co-occurrence based on their genomic linear distance, assuming their homogeneous distribution in their chromosome volume (Beagrie et al. 2017; Beagrie et al. 2023). For all possible three-way locus contacts, SLICE calculates the probability of each triplet interaction (*Pi3*). The top 2% triplets, ranked according to their average probability of interaction, were selected for further analysis as the most prominent triplet interactions. SLICE identified approx. 4×10^6^ and 5×10^6^ significant (top 2%) three-way interactions in ESCs and in DNs, respectively (**GSE304996**).

### ESC and DNs have specific chromatin organization

To begin studying cell-type specific chromatin interactions in ESCs and DNs, we visualized GAM contact matrices centered on the *Sox2* gene locus (**Figure 1C**). *Sox2* is expressed in both ESCs and DNs, where it plays different roles. In ESCs, it acts as a pluripotency factor essential for cell survival (Li et al. 2014; Zhou et al. 2014), while in neurons it is important for neuronal differentiation (Zhang and Cui 2014). In ESCs, approx. 90% of *Sox2* expression is controlled by a SE located 100 kb upstream of its promoter (Li et al. 2014; Zhou et al. 2014). *Sox2* forms significant pairwise (*Pi2)* interactions with the upstream ESC-specific SE only in ESCs but not in DNs (**Figure 1C**). GAM captures significant pair-wise interactions with more distal regions upstream and downstream of *Sox2*, both with other genes and with intergenic regions. In DNs, novel and more abundant *Sox2* interactions are detected, especially with the most distal regions. Distal chromatin interactions at the *Sox2* locus, beyond the TAD level, were also identified by high sequencing depth Hi-C using *in-vitro* differentiated pyramidal glutamatergic neurons (Bonev et al. 2017; **Supplemental Figure S1G**). Using the top 2% most significant *Pi3* interactions obtained by SLICE, we find that the 150 kb genomic window containing *Sox2* establishes many three-way interactions in both cell types, with a larger number of genomic partners in DNs. These results highlight extensive differences in chromatin organisation between ESCs and DNs at both local and distal levels of chromatin organisation that involve pairwise and three-way chromatin interactions (**Figure 1C**).

### Mapping differential gene expression and chromatin accessibility in ESCs and DNs

To study how the structural reorganisation of chromatin contacts in ESCs and DNs relates with changes in gene expression and chromatin remodelling, we measured gene expression using total RNA-seq and chromatin accessibility using ATAC-seq. We collected total RNA-seq from ESCs and DNs (four biological replicates; **Supplemental Figure S2A; GSE305150**), and detected 8,942 genes are expressed in both cell types, and 909 and 3998 genes expressed only in ESCs or DNs, respectively (TPM ≥ 1 in > 2 biological replicates; **Supplemental Figure S2B**). Differentially expressed genes were identified using DESeq2 (Love et al. 2014), applying stringent criteria (adjusted p-value < 0.05, log2(Fold Change) > |2|, |dTPM| > 1; **Figure 2A; Supplemental Table S5**).

**Figure 2.**
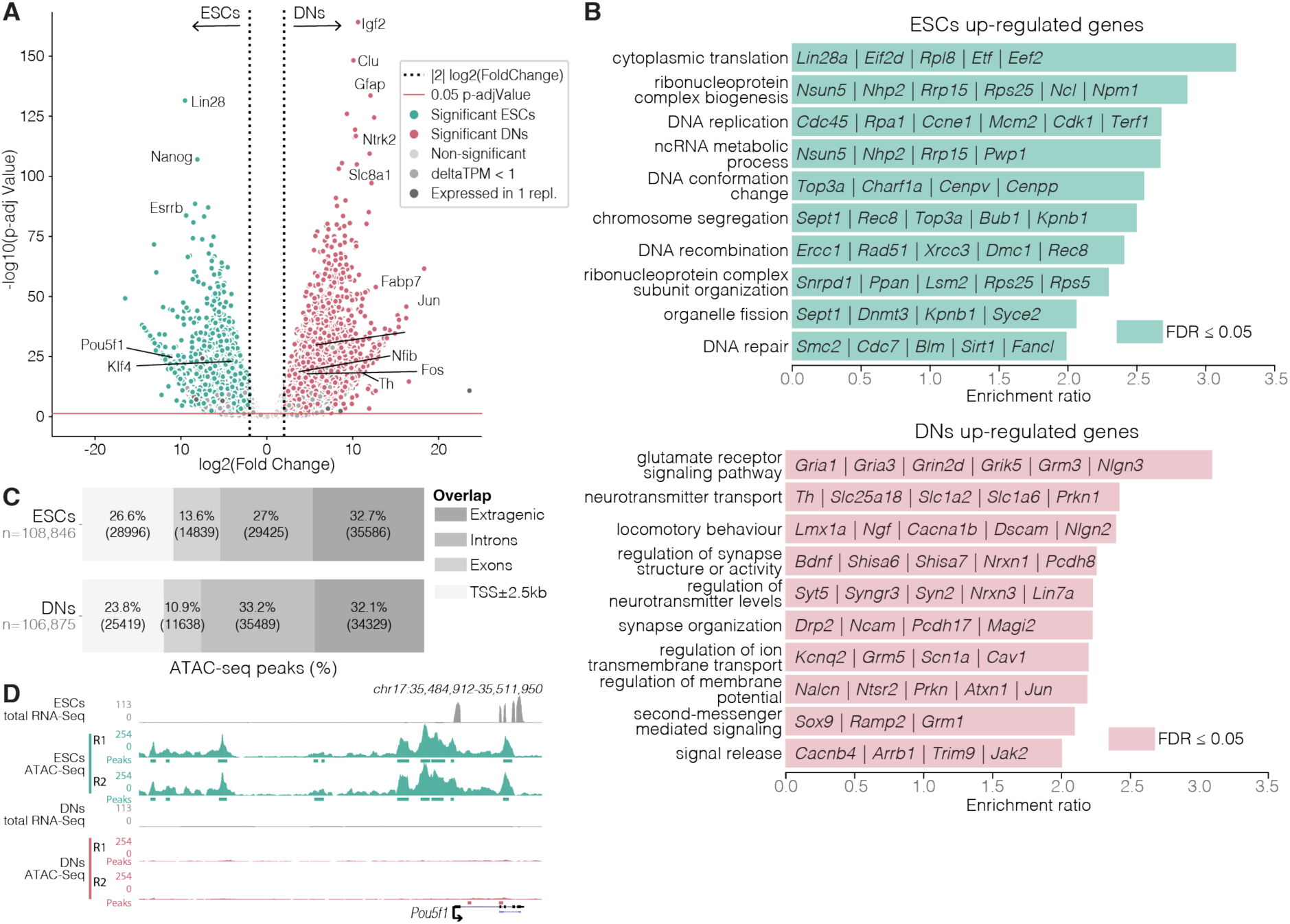
Gene expression and chromatin accessibility in mouse embryonic stem cells (ESCs) and dopaminergic neurons (DNs). (**A**) Volcano plot showing the distribution of the log2(Fold Change) and log10(adjusted p-value) from DESeq2 applied in four replicates of mouse embryonic stem cells (ESCs) and dopaminergic neurons (DNs) using total RNA-Seq samples. Dotted red and black lines represent the significance thresholds applied (|log2(Fold Change)| > 2, adjusted p-value < 0.05). Genes were considered differentially expressed if they passed the significance thresholds, were expressed ( ≥ 1 TPM) in more than two replicates, and had a TPM difference higher than 1 TPM (|dTPM| > 1). Arrows indicate examples of differentially expressed genes. (**B**) Barplot showing the Gene Ontology (GO) enrichment analysis for the genes classified as up-regulated in ESCs or in DNs. GO terms are considered significantly enriched when the False discovery rate (FDR) ≤ 0.05. Examples of genes from each GO are highlighted. (**C**) Stacked bar plot showing the percent of ATAC-Seq peaks classified based on their overlap with extragenic regions, introns, exons and promoters (TSS±2.5 kb) of known genes. (**D**) Examples of tracks for expression (total RNA-Seq) and chromatin accessibility (ATAC-Seq) for two biological replicates, in mouse embryonic stem cells (ESCs) and dopaminergic neurons (DNs). *Pou5f1* (POU Class 5 Homeobox 1) is a pluripotency marker for ESCs.

Amongst the 2,476 genes up-regulated in ESCs, we find pluripotency factors such as *Klf4*, *Nanog* or *Pou5f1*, while the 3,329 genes up-regulated in DNs include neuronal activity genes *Fos* and *Jun*, and dopaminergic markers *Th* and *Fabp7* (**Figure 2A**). Gene ontology (GO) analysis showed that genes up-regulated in ESCs are significantly enriched for terms associated with metabolic processes, DNA repair or processes associated with cell division, such as DNA replication or chromosome segregation (**Figure 2B**, top panel; **Supplemental Table S6**). Genes up-regulated in DNs show enrichment for terms associated with neuronal-specific functions, such as neurotransmitter transport, synapse organisation or regulation of membrane potential (**Figure 2B**, bottom panel; **Supplemental Table S6**). Overall, ESCs and DNs showed cell-type specific transcriptional profiles fine-tuned to preserve and accommodate specialised cellular functions.

To profile chromatin accessibility in ESCs and DNs, we applied ATAC-Seq in two biological replicates (**Supplemental Figure S2C; GSE304720**). We identified robust ATAC peaks shared between replicates (108,846 and 106,875 peaks in ESCs and DNs, respectively), and classified them based on their overlap with extragenic regions, annotated promoters or coding regions (Harabula et al. 2025). Most ATAC peaks were present in extragenic or intronic regions in both cell types (**Figure 2C**). We confirmed that some of the extragenic peaks overlap regions with known regulatory functions. For example, the expression of the pluripotency factor *Pou5f1* in ESCs, coincides with considerable ESC-specific chromatin accessibility upstream and across the gene (**Figure 2D**). These results show that most changes in chromatin accessibility occur in non-coding genomic regions potentially involving regulatory elements such as SEs.

### Super-enhancers detected in ESCs and DNs are highly cell-type specific

To study the role of SEs in chromatin organisation in ESCs and DNs, we next identified active enhancers based on ATAC-seq peaks which are positive for H3K27ac and H3K4me1 chromatin occupancy (Hnisz et al. 2013) and negative for H3K27me3 (Rada-Iglesias et al. 2011; Cruz-Molina et al. 2017; Harabula et al. 2025; **Supplemental Figure S3A**). The chromatin occupancy of H3K27ac and H3K4me1 was profiled genome-wide using ChIP-Seq in matching ESC and DN populations. H3K27me3 ChIP-Seq datasets collected from the same system were obtained from publicly available resources (Ferrai et al. 2017). A total of 8,519 and 9,699 putative active enhancers which do not overlap promoters (±2.5 kb) or exons were identified in ESCs in DNs, respectively (**Supplemental Figure S3B**).

Next, we identified SEs in ESCs and DNs by applying the ROSE pipeline (Lovén et al. 2013; Whyte et al. 2013). We uncovered 906 putative SEs in ESCs and 628 in DNs (**Figure 3A; Supplemental Table S7**). The SEs identified in ESCs include most SEs that were previously described based on histone modifications and TF occupancy (**Supplemental Figure S3D**; Whyte et al. 2013). The average length of ESC and DN SEs was 35 and 52 kb, respectively (**Supplemental Figure S3E**), consistent with previous studies (Hnisz et al. 2013). Taken together, these results show that DNs have fewer but longer SEs than ESCs.

**Figure 3.**
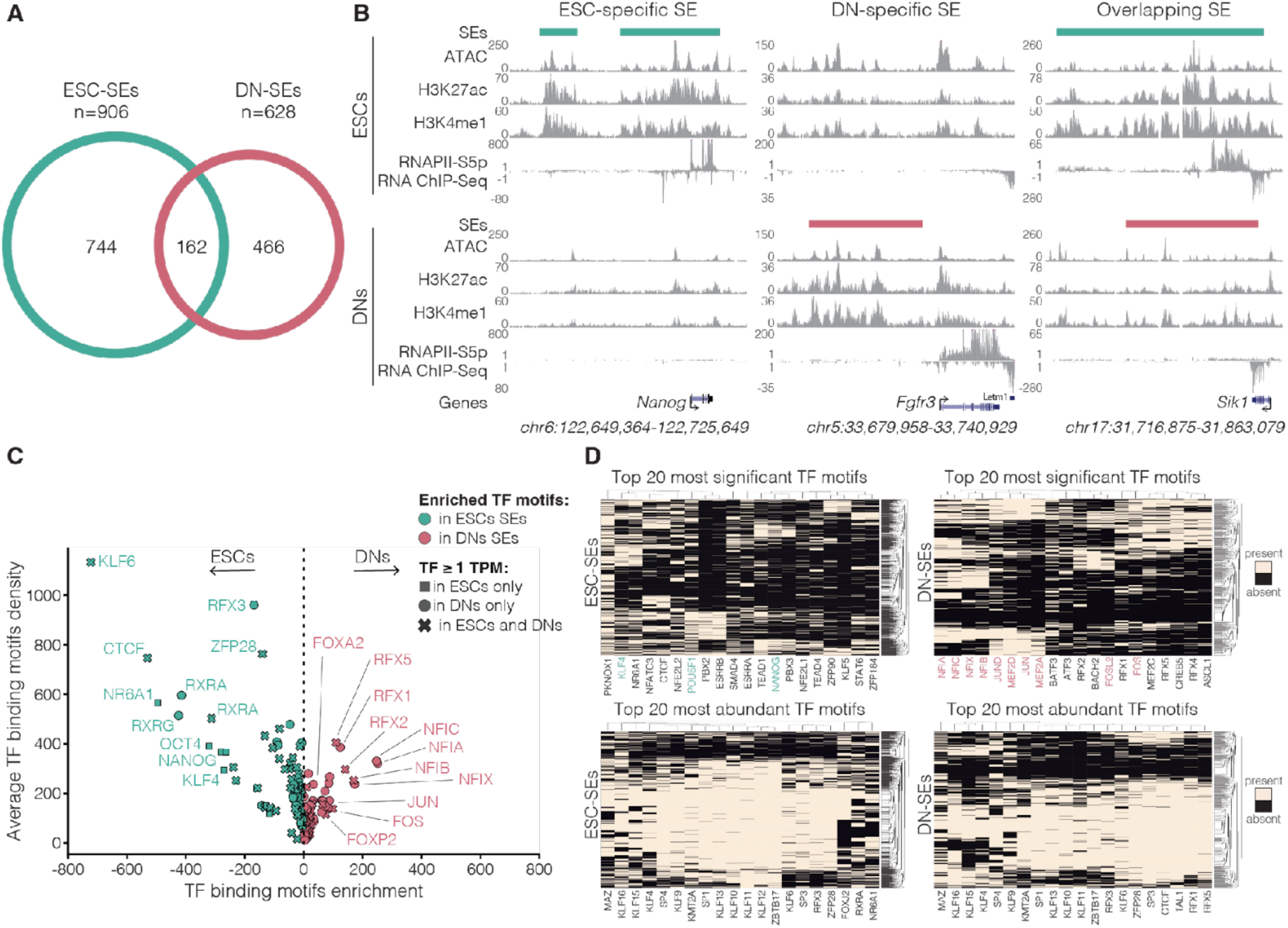
Cell-type specific super-enhancers (SEs) in mouse embryonic stem cells (ESCs) and dopaminergic neurons (DNs). (**A**) Number of cell-type-specific and shared SEs in ESCs and DNs. Shared SEs overlap by at least 1 bp. (**B**) Examples of UCSC genome-browser tracks in ESCs and DNs for ESC-specific, DN-specific and common (overlapping) SEs. Tracks plotted for SEs, chromatin accessibility (ATAC-Seq), ChIP-Seq for histone H3 acetylated on lysine K27 (H3K27ac ChIP-Seq) and for histone H3 mono-methylated on lysine K4 (H3K4me1 ChIP-Seq), and nascent RNA (RNAPII-S5p RNA-ChIP-Seq). RefSeq gene tracks for *Nanog*, *Fgfr3* (Fibroblast growth factor receptor 3) and *Sik1* (Salt Inducible Kinase 1). (**C**) Volcano plot showing the differential transcription factor (Dunham et al.) motifs found significantly enriched in ESC-specific or DN-specific SEs. Expressed TFs have ≥ 1 TPM. Differentially enriched TF motifs were identified using an independent t-test. Significant motifs were defined as p < 0.05. (**D**) Heatmap showing the absence/presence of the top-20 most significantly enriched (top row) and top 20 most abundant (bottom row) motifs of expressed TFs found in ESC and DN SEs. Examples of interesting TFs are highlighted with colours.

Most SEs identified in ESCs and DNs were cell-type specific (**Figure 3A**). For example, two ESC-specific SEs are present at the *Nanog* locus, expressed only in ESCs (**Figure 3B**, left panel). One DN-specific SE is detected upstream the *Fgfr3* locus (**Figure 3B**, centre panel), a gene which is significantly upregulated in DNs (**Supplemental Table S5**), and has crucial roles in DN development and survival (Timmer et al. 2007). A small group of 162 SEs are present in both ESCs and DNs, and may regulate genes expressed in both cell types, such as *Sik1* (**Figure 3B**, right panel), which encodes a kinase with a wide range of functions in cellular homeostasis (Sun et al. 2020).

We characterized the genomic position of the SEs detected here, and found that they are present across all mouse chromosomes with some chromosomes showing higher SE numbers in ESCs (e.g. chromosomes 3, 10 and 13; **Supplemental Figure S3F**). More than 50% of SEs overlap with expressed genes in both ESCs and DNs, and fewer overlapped DEGs in ESCs (20%) or DNs (25%; **Supplemental Figure S3H; Supplemental Table S7**). The co-occurrence of SEs and expressed genes in the linear genome suggests that some genes may serve as regulatory hubs. Overlap between coding regions of expressed genes and H3K27ac positive regions has been previously reported in neurons (Zhao et al. 2018).

### ESC– and DN-specific SEs are enriched for cell-type-specific TFs

Next, we investigated more deeply the properties and specificity of SEs in ESCs and DNs. SEs were previously shown to be enriched for cell-type-specific TFs (Hnisz et al. 2013; Whyte et al. 2013). To discover enriched TF binding motifs in the ESC and DN SEs, we developed a pipeline to identify significantly enriched TF binding motifs (considering only TFs expressed in either cell type) within accessible chromatin regions (**Supplemental Figure S3G**).

Our approach identified 88 and 73 TF motifs enriched in ESC– and DN-specific SEs, respectively (**Figure 3C**; **Supplementary Figure S3I; Supplemental Table S8**). ESC-specific SEs were enriched for motifs of master pluripotency factors KLF4, OCT4 and NANOG (**Figure 3C**; Whyte et al. 2013), and pluripotency maintenance factor ZIC (Lim et al. 2007). In contrast, DN-specific SEs were enriched for motifs of TFs associated with neuronal differentiation, such as NFIs (Harris et al. 2015) and RFXs (Harris et al. 2021), and neuronal signalling, including FOS and JUN (Chung 2015). SEs shared between ESCs and DNs also showed enrichment of cell-type specific expressed TFs (**Supplemental Figure S3J; Supplemental Table S8**), suggesting that there is specificity in the TFs that potentially bind and regulate the function of common regulatory regions.

SEs can be bound by both homotypic and heterotypic clusters of TFs which dictate the properties and regulatory functions of the SEs (Wang et al. 2025). To determine which TF combinations are found in ESC and DN SEs, we looked at the most significantly enriched TFs and the most abundant TFs (**Figure 3D; Supplemental Table S8**). Different combinations of TF motifs were enriched in ESC SEs than in DN SEs, with the largest variation in the most significantly enriched motifs, whilst TF motifs that were most abundant were shared between the two cell types. In ESC SEs, the most significant motif combinations were formed of TFs important for pluripotency, such as KLF4, OCT4 or NANOG. In contrast, the most significant motif combinations in DNs were for TFs important for neuronal development, such as NFI TFs, or with neuronal activity, such as FOS and JUN. The most abundant motifs in ESC– and DN-specific SEs were putative binding sites for universal TFs, such as MAZ, SP1 or CTCF. These findings suggest that SE activity is regulated through recruitment of distinct combinations of TFs for both shared SEs and cell-type-specific SEs, prompting us to investigate their architectural and regulatory roles in more detail.

### SEs and DEGs form pairwise interactions that span megabases in both ESC and DNs

To investigate the architectural roles of SEs in ESCs and DNs, we searched for the connections between SEs and DEGs in the most significant (top 5%) SLICE pairwise (*Pi2*) chromatin interactions at 5 kb resolution (**Supplemental Figure S4A; GSE304996**). We found that most SEs form pairwise interactions with expressed genes, but remarkably approximately 80% of ESC SEs and 50% of DN SEs interact with DEGs (**Supplemental Figure S4B**).

Next, we searched for the most represented types of contacts established by SEs and/or DEGs. Most pairwise SLICE interactions are established between genes upregulated in ESCs or DNs, followed by interactions between the upregulated genes and SEs, and by SE interactions with other SEs (**Figure 4A**). Considerably more prominent interactions are detected in DNs than ESCs across all compared classes of contacts. For example, DN-upregulated genes establish 5.5×10^6^ prominent interactions in DNs, compared with 0.2×10^6^ interactions involving ESC-upregulated genes in ESCs. Furthermore, most upregulated genes interact at least once with SEs in the corresponding cell type (> 80%; **Figure 4B**). SE interactions are not limited to DEGs, and also involve a high proportion of expressed and non-expressed genes (**Supplemental Figure S4C**).

**Figure 4.**
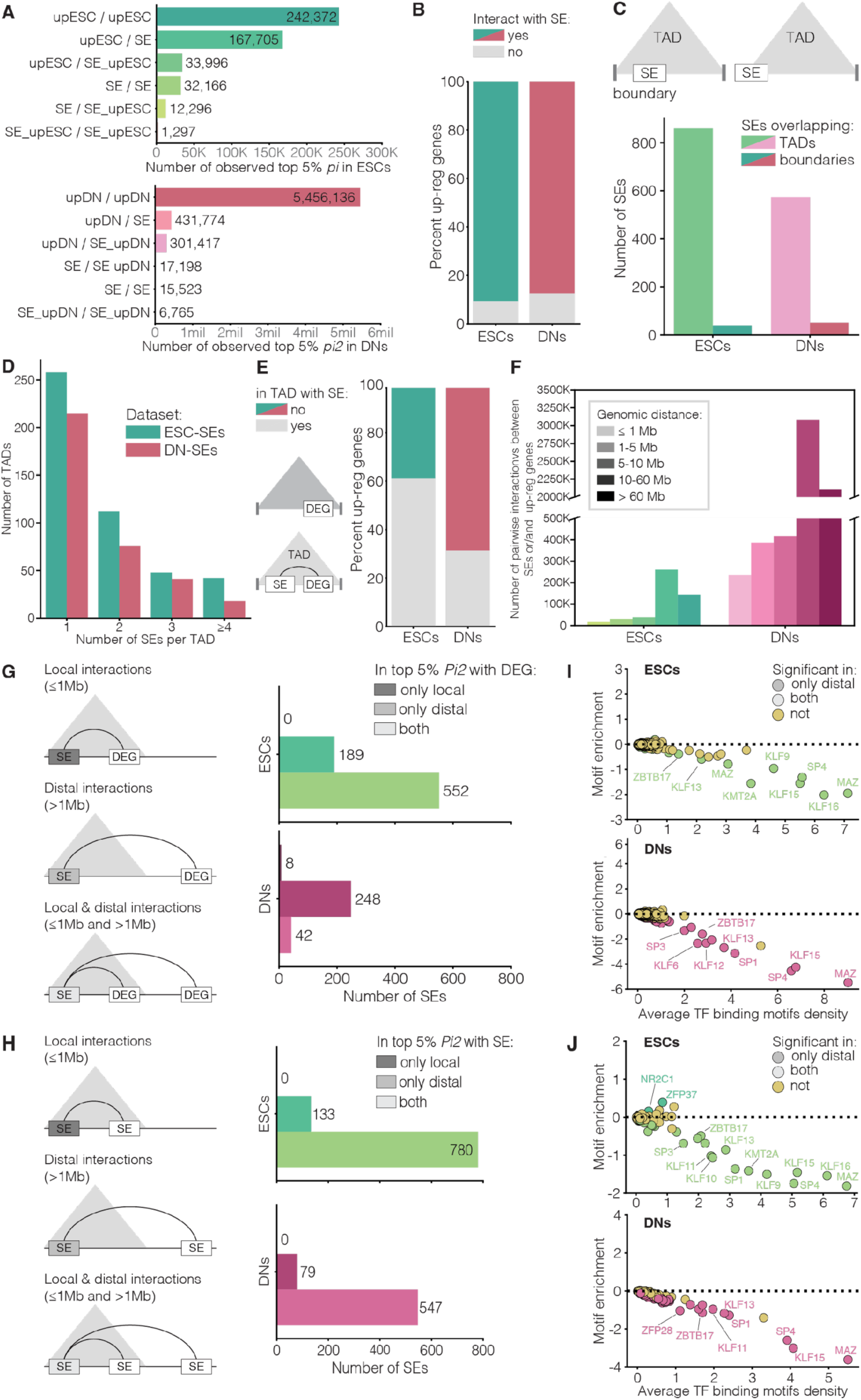
Super-Enhancers (SEs) are involved in mega-base range pairwise interactions with other SEs and differentially expressed genes (DEGs) in mouse embryonic stem cells (ESCs) and dopaminergic neurons (DNs). (**A**) Barplot showing the number of top 5% SLICE pairwise (*Pi2*) interactions, in ESCs and DNs, between 5 kb windows classified based on their overlap with upregulated genes (up-ESCs/up-DNs), SEs (SE) or both (SE-up) in ESCs (top panel) or DNs (bottom panel). (**B**) Stacked barplot showing the proportion of upregulated genes in ESCs and DNs found in top 5% SLICE interactions with SEs. (**C**) Barplot showing the number of SEs in ESCs and DNs present within topologically associating domains (TADs) or that overlap TAD boundaries. (**D**) Barplot showing the number of SEs per TAD in ESCs and DNs. (**E**) Stacked barplot showing the proportion of upregulated genes in ESCs and DNs found in the same TAD with at least one SE. (**F**) Barplot showing the number of top 5% SLICE pairwise interactions between SEs and upregulated genes in ESCs and DNs, at different genomic distances. (**G**) Barplot showing the number of top 5% SLICE pairwise interactions between SEs and genes upregulated in ESCs (top panel) or DNs (bottom panel), classified based on their genomic distance into local, distal or both local and distal interactions. (**H**) Bland-Altman plot showing the enrichment in expressed TF motifs in ESCs SEs (top panel) and DNs SEs (bottom panel), colour coded based on whether SEs form distal, or distal-and-local top 5% SLICE pairwise interactions with upregulated genes. Differential motif presence was assessed with an independent sample t-test. Analysis includes only motifs of expressed TFs (≥ 1 TPM ESCs U DNs). (**I**) Barplot showing the number of SEs in top 5% SLICE pairwise interactions with SEs in ESCs (top panel) and DNs (bottom panel) at different genomic distances, classified into local, distal or both local-and-distal interactions. (**J**) Bland-Altman plot showing the enrichment in expressed TF motifs in ESCs SEs (top panel) and DNs SEs (bottom panel), colour coded based on whether the gene forms distal or distal-and-local top 5% SLICE pairwise interactions with an SE. Differential motif presence was assessed with an independent sample t-test. Analysis includes only motifs of expressed TFs (≥ 1 TPM ESCs U DNs) selected from the HOCOMOCO motif database.

Current reports show that SEs can regulate gene expression via short-range chromatin interactions that occur within TADs (Jia et al. 2019). To investigate the extent of SE interactions within and between TADs with other SEs and DEGs, we called TADs using the insulation score method at 50 kb resolution (see **Methods**; **Supplemental Table S9**) (Winick-Ng et al. 2021). In both ESCs and DNs, most SEs are found within TADs and do not overlap with their boundaries (**Figure 4C**), and most TADs contain one or two SEs (**Figure 4D**). Interestingly, 40% of upregulated genes in ESCs and approx. 65% of upregulated genes in DNs are found in TADs devoid of SEs (**Figure 4E**), suggesting that differentially expressed genes are either not regulated by SEs or establish interactions with SEs that cross TAD boundaries. To investigate this hypothesis, we counted the number of pairwise interactions between SEs and upregulated genes at different genomic distances. We found many interactions between SEs and DEGs, in both ESCs and DNs, but they most often occur between 10 and 60 Mb (**Figure 4F**). We also investigated SE-SE interactions and found that they also preferably occur across large genomic distances of 10 and 60 Mb (**Supplemental Figure S4D**). After classifying SEs according to the distance of their pairwise interactions with DEGs, we find very few SEs involved only in local interactions (<1Mb), while many SEs establish distal-only (>1Mb) or local-and-distal (≤ 1Mb and >1Mb) interactions with upregulated genes (**Figure 4G**). The same behaviour is observed for SE-SE interactions (**Figure 4H**), and for DEG-SE interactions (**Supplemental Figure S4E**). These results suggest that SEs form an extensive network of pairwise regulatory interactions with DEGs and other SEs spanning large genomic distances beyond the TAD level.

### MAZ, KLF and SP TF motif families are enriched in SE-DEG interactions

To determine if specific TFs distinguish SEs involved only in distal interactions with DEGs, but not in SEs involved in both local and distal interactions, we performed differential TF binding motif analysis using an independent sample t-test for each cell type. We found significant enrichments of motifs from the MAZ, KLF and SP TF families in ESC SEs and DN SEs that are involved in both local and distal interactions with DEGs (**Figure 4I; Supplemental Table S10**), even though most SEs are cell-type specific. In contrast, SEs involved only in distal interactions do not have specific TF motif enrichments, suggesting that they are not generally associated with a single TF. Interestingly, TFs from families such as MAZ, KLF and SP are also identified as significantly enriched in the SEs or DEGs that interact with other SEs both locally and distally, compared to SEs that establish only distal interactions (**Figure 4J**; **Supplemental Figure S4F**). In DNs, the DEGs involved only in distal interactions with other SEs are enriched for DN-specific TFs, such as the FOX family of TFs (**Supplemental Figure S4F** bottom panel**; Supplemental Table S10**). These results suggest that primary TFs may be important for interactions between SEs and/or DEGs that are both local and distal, while distal-only interactions can involve cell-type-specific sets of TFs.

### Complex networks of SE interactions in both ESCs and DNs

Genome-wide studies indicate that complex chromatin interactions that assemble many genomic regions may play a role in modulating gene expression and may be important for understanding disease pathomechanisms (Beagrie et al. 2017; Deshpande et al. 2022). The observation that SEs are preferentially involved in both local and distal interactions with other SEs raises the question of whether these interactions are simultaneous or occur in different cells. To identify three-way interactions between SEs and explore their properties, we used the significant three-way interaction probabilities (*Pi3*) calculated using SLICE (Beagrie et al. 2017; Beagrie et al. 2023) at 150 kb resolution. First, we asked whether three-way interactions are enriched for SEs and highly transcribed regions, as previously shown in ESCs at a lower genomic resolution of 1Mb (Beagrie et al. 2017). We classified the 150 kb windows first according to SE presence, and for the remaining windows without SEs according to their transcriptional activity (low, medium-low, medium-high and high; **Supplemental Figure S5A; Supplemental Table S11**). Transcriptional activity of genomic windows was quantified from chromatin-associated RNA transcripts pulled down in association with RNAPII-S5p, by RNA ChIP-Seq using antibodies against RNAPII-S5p that detect RNAPII at gene promoters and throughout the coding region (Brookes et al. 2012). SE-containing windows were typically highly transcribed in both ESCs and DNs, with 82% or 73% being ranked as high transcription, respectively.

Next, we classified the prominent triplet interactions according to their window classifications in ESC and DNs (**Supplemental Table S11**), and tested which combinations were significantly enriched over random expectation genome-wide, estimated from 1000 random linear, genomic distance-preserving permutations of the triplet window coordinates. The most prominent triplet interactions in both cell types involve SE/SE/SE interactions, followed by interactions between SEs and highly transcribed genomic windows (**Supplemental Figure S5B**). These observations extend previous findings of three-way interactions between TADs that contained SEs (Beagrie et al. 2017), which we now detect with more specificity, higher resolution, and enrichment. For example, we find 12,025 prominent SE/SE/SE interactions in ESCs compared to 200 described in prior work (Beagrie et al. 2017). Triplets involving high-transcription windows were also highly abundant (244,374), occurring often in combination with SE-containing windows. Conversely, most depleted three-way interactions contained low or medium-low transcription genomic windows, and rarely contained SE windows, but no depletion was significant (i.e., p-value >= 0.05). Further, we found a smaller number of SE/SE/SE interactions in DNs (2,675), compared with ESCs (12,025), as well as three-way interactions between highly transcribed windows (127,361 and 244,374 in DNs and ESCs, respectively).

### ESC– and DN-specific SEs preferentially establish three-way interactions

Since SEs can be cell-type specific or common to ESCs and DNs, we asked whether the most prominent triplet interactions were enriched for cell-type-specific SEs or common SEs. We classified SEs as either common, ESC-specific or DN-specific, and performed a permutation analysis to identify the most prominent three-way chromatin interactions. Remarkably, in both ESCs and DNs, the prominent triplet interactions were enriched for cell-type-specific SEs (**Figure 5A**), suggesting that cell-type-specific SEs are especially important for long-range complex interactions which in turn may play a role in regulation of cell-type-specific gene expression. At the top of the significant enrichments were also triplets between common and cell-type-specific SEs, or highly transcribed windows with cell-type-specific SEs, emphasizing the specificity and complexity of the regulatory networks involving three-way interactions.

**Figure 5.**
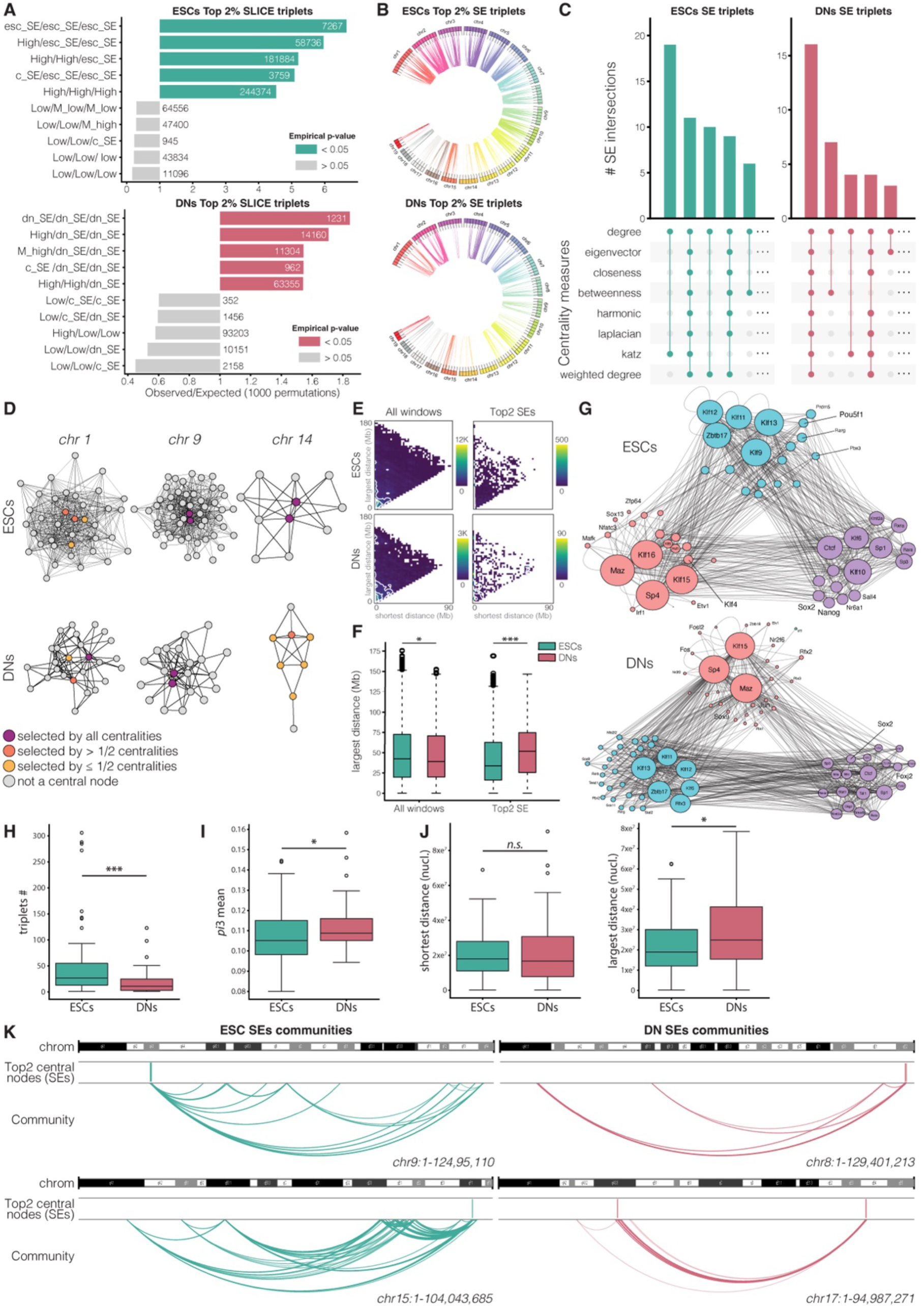
Cell-type specific complex chromatin interactions between Super-Enhancers (SEs) in mouse embryonic stem cells (ESCs) and dopaminergic neurons (DNs). (**A**) Barplots showing the top-5 and bottom-5 enrichment analysis results for top 2% significant three-way interactions detected by SLICE in GAM data from ESCs (top) and DNs (bottom), at 150 kb resolution. For each cell type, windows overlapping SEs were classified into common SEs (c_SE; i.e. overlapped shared SEs) or ESCs-specific SEs (esc_SE) and DNs-specific SEs (dn_SEs). The length of the bar represents the observed number of triplets divided by the expected number obtained from 1000 permutations. Bars are coloured based on the permutation test empirical p-value. Numbers indicate the observed interactions detected between 150 kb windows in each category. (**B**) Circos plots showing the top 2% three-way SLICE interactions between SEs across all mouse chromosomes in ESCs (top) and DNs (bottom). (**C**) UpSet plots showing the top SEs classified based on their centrality measures. Eight centrality measures were used to select the top-2 central SE nodes in each SE interaction network for each cell type. (**D**) Examples of SE networks identified in different chromosomes in ESCs and DNs. SE nodes are coloured based on how many centrality measures selected the particular SE. (**E**) Density plots showing the distances between two nodes of a SE triplet. Shortest and longest distances are on the x and y axes, respectively. Distances are displayed for all possible triplets, and for triplets present in the SE network that contain a node selected by at least one centrality measure. (**F**) Boxplots showing a comparison between ESCs and DNs of largest distances identified between SEs in triplets. Comparison done using a t-test. *p-value < 0.05, **p-value< 0.01, ***p-value<0.001, n.s.non-significant. (**G**) Networks of transcription factors for which motifs were identified in SEs involved in three-way contacts. Leiden community detection was used to identify strong groups of interacting SEs. Colours indicate the three communities of motifs identified in ESCs and DNs. (**H**, **I**, **J**) Boxplots show comparisons between ESC and DN communities of SEs, for triplet counts (H), mean interaction strength (I), and interaction distances (J). Comparisons performed using the two-sided Wilcoxon rank-sum test. *p-value < 0.05, **p-value< 0.01, ***p-value<0.001, n.s.non-significant. (**K**) WashU browser showing examples of top SE communities in ESCs and DNs ranked based on the number of top-2 central nodes identified in the community normalised by the total number of SEs identified per chromosome.

### Centrality analysis identifies key SEs in chromatin interaction networks

SEs are a major component of three-way chromatin interactions in both ESCs and DNs, irrespective of whether they are shared or common. To further explore the regulatory importance of SEs, we modelled the SE-SE-SE triplet interactions as a weighted network, where a node represents an SE and a weighted edge represents the median of *Pi3* values of all interactions involving the node. SE-SE-SE triplets are found in all chromosomes, except for chromosome 13 in DNs (**Figure 5B**). To explore the regulatory importance of each SE in the interaction network, we calculated eight centrality measures: degree, eigenvector (Bonacich 1972), closeness (Freeman 1978), betweenness (Brandes 2001), harmonic (Boldi and Vigna 2014), Laplacian (Qi et al. 2012), Katz (Katz 1953) and weighted degree (Zuo et al. 2012; S**upplemental Figure S5C**). Remarkably, a subset of SEs found in most chromosomes ranked the highest across all centrality measures (11 SEs in ESCs and 16 SEs in DNs; **Figure 5C**; all combinations shown in **Supplemental S5D**; **Supplemental Tables S12-S13**), suggesting that these SEs may function as major regulatory hubs. These high-centrality SEs exhibited high local connectivity (degree centrality) and strong global influence (eigenvector and Katz centralities), whilst also occupying key positions in the network (betweenness centrality). Their high closeness and harmonic centrality scores further indicate that they may play important roles as relays of regulatory information in the networks. Interestingly, several SEs exhibited more nuanced centrality, ranking highly only in a subset of centrality measures. For instance, some SEs are characterised by high scores in local connectivity (e.g., high degree centrality) or efficiency of access (e.g., high closeness centrality), but had low scores for influence-based measures such as eigenvector centrality. Amongst the highest-ranking SEs, 75 are ESC specific and 50 are DN specific, while only 3 and 12, are common SEs, respectively (**Supplemental Table S13**). Taken together, these results suggest that high-ranking SEs may act as specialised relays for regulatory information, rather than universal regulatory hubs. The diversity in the centrality profiles likely reflects the functional heterogeneity of SEs within the chromosomal networks of triplet chromatin interactions.

To visualise the positions of high-centrality SEs within the SE triplet interaction networks, we constructed interaction networks for individual chromosomes, where nodes represent SEs coloured based on their centrality profiles: SEs scoring high across all centrality measures, and SEs scoring high in either more than half of the centrality measures or fewer than half of the centrality measures or none of the centrality measures (**Figure 5D**). These visualizations reveal distinct patterns of SE contributions across chromosomes, and specific and common properties of the SE-SE-SE triplet interaction networks between ESCs and DNs (**Supplemental Figure S6A,B**). In some cases, such as chromosome 9, two SEs in each cell type have high scores across all centrality measures, while being surrounded by SEs with low or no centrality signals. In other cases, such as chromosome 1 in ESCs, there are no SEs with high centrality scores across all measures, but SEs with intermediate centrality profiles dominate. These results suggest that SEs have non-uniform roles within the networks and in each cell type.

To investigate whether SEs having high centralities establish triplet interactions with preferred genomic length, we compared the largest and shortest genomic distances between SEs in triplets containing at least one highest-ranked SE with SEs in all triplets (**Figure 5E**). Considering the two highest centrality SEs in each chromosome, we found that the genomic lengths of their triplets are preferentially shorter in ESCs compared to DNs. The largest distances of the top 2 centrality triplets are typically around 40 or 90 Mb in ESCs and DNs, respectively (**Figure 5F**). In contrast, the largest genomic distances of SE triplets involving the bottom 2 centrality SEs are not significantly different between ESCs and DNs (**Supplemental Figure S7A-C**). These results suggest that cell-type-specific enhancer networks may be organized on different genomic scales, potentially reflecting distinct regulatory strategies in post-mitotic DNs, which may be important for preserving cell type identity over large periods of time.

### SEs in networks are enriched for distinct communities of transcription factors

To discover putative activities and relationships of TFs involved in the SE chromatin interaction networks, we used the TF binding motifs present within the participating SEs to create a network wherein each node represents a TF and each edge represents a TF pair interaction implied by SE interactions. We detected the most interconnected communities in the TF network using the Leiden algorithm (Traag et al. 2019). We visualised the resulting TF communities using a network wherein node colour represents a Leiden-assigned TF community, the node size represents the cumulative abundance of a TF across all SEs, and an edge weight encodes the frequency of a specific TF interaction (inferred from the SE interactions). We find three major TF communities in both ESCs and DNs which contain cell-type specific TFs that are expressed in similar levels in both cell types (**Figure 5G**). The TF communities identified likely reflect distinct regulatory features of the SE networks in both cell types. For instance, we find a TF community which includes MAZ, KLF15 and SP4 as the most abundant in SEs involved in SE-SE-SE contacts in both ESCs and DNs, but also cell-type-specific TF community members, such as KLF4 in ESCs, and SOX9, JUN and FOS in DNs. These results further suggest that different groups of SEs may be specialised in controlling specific biological functions or chromatin states.

### Modular organization of SE interaction networks revealed by community detection

We investigated whether the SEs in the triplet-contact network are organised in communities containing SEs that interact more with each other than with other SEs. We identified several SE communities (an average of 4 communities per chromosome in ESCs and 3 communities per chromosome in DNs; **Supplemental Table S14**). SE communities have a significantly higher number of SE triplets in ESCs compared to DNs (**Figure 5H**), but the mean probability of interaction (*Pi3*) within these communities is higher in DNs than ESCs (**Figure 5I**). The genomic distances between interacting SEs in DN communities are significantly larger in comparison to ESCs (**Figure 5J**). We visualised SE communities using the WashU genome browser (Li et al. 2019), and found that the SE communities can span whole chromosomes, in various configurations (**Figure 5K**; **Supplemental Tables S15-S16**). Interestingly, in both ESCs and DNs, some communities include SEs that are ranked as the most central nodes in the network analyses, suggesting that they may have specific roles in different sections of the networks, and may act as regulatory hubs (**Supplemental Table S14**). These results show that SE communities have different properties between dividing ESCs and post-mitotic DNs, and provide a resource for prioritizing SEs with different importance in 3D genome organisation and function.

## Discussion

In this study, we investigated the contributions of SEs to chromatin architecture in mouse ESCs and *in vitro* DNs. By integrating high-resolution GAM data with gene expression, chromatin accessibility and histone modifications, we identified cell-type specific SEs and revealed their extensive involvement in long-range and complex three-way chromatin interactions. This work expands our understanding of SE contributions to the architectural organisation of the genome in specific cell types, by showing that SEs establish complex regulatory networks of cell-type specific interactions with other SEs and genes with specialized cellular functions.

### SEs form extensive megabase-range chromatin interactions in ESCs and DNs

Our data show that most SEs are highly cell-type specific, consistent with their well-established role in driving lineage-specific gene expression programs (Whyte et al. 2013). ESC-specific SEs are enriched for binding motifs of pluripotency-associated TFs, while DN-specific SEs are marked by motifs of TFs involved in neuronal development and activity. SEs are composed of regions with different regulatory capacities (Blayney et al. 2023), as reflected by the presence of different TF binding motifs. Different combinations of cell-type specific TFs may also facilitate the complexity of the long-range pairwise and three-way interactions established by SEs and contribute to their cell-type specific regulatory functions. Strikingly, many complex interactions involving SEs and DEGs span tens of megabases in both ESCs and in neurons, connecting distant TADs and suggesting a broader functional range for SE activity than previously appreciated.

### Complex SE chromatin interactions are highly cell-type specific

A key finding from our study is that cell-type-specific SEs are found in complex chromatin interactions in both ESCs and DNs. DNs have fewer SEs and fewer SE-SE-SE interactions than ESCs, but the SE triplets present in DNs establish more frequent interactions that occur over significantly larger genomic distances compared to ESCs. These results are consistent with a more compartmentalized and specialized regulatory architecture in terminally differentiated, post-mitotic cells, as previously suggested (Fraser et al. 2015; Bonev et al. 2017; Winick-Ng et al. 2021). The increased genomic distances spanned by interactions involving network-central SEs indicate a reorganization of the regulatory landscape during neuronal differentiation toward long-range coordination of transcriptional hubs. Rewiring of chromatin interactions during neuronal differentiation has been previously observed during neuronal lineage commitment (Fraser et al. 2015) and is associated with the reconfiguration of Polycomb-associated gene networks as they transition from a poised state to an actively transcribed state (Bonev et al. 2017).

TF motif analyses revealed that distinct SE modules are enriched for different combinations of TF motifs, including both ubiquitous and cell-type-specific regulators. This supports the notion that SEs are not homogeneous regulatory units, but rather serve as multivalent scaffolds that integrate different TF combinations to drive specific gene expression programs. The recruitment of diverse TF communities to SE hubs likely enhances the specificity and robustness of transcriptional control in both pluripotent and terminally differentiated states.

Through network community detection, we identified chromosomal sub-networks of SE interactions that may reflect their functional compartmentalization. These SE communities differ between ESCs and DNs in size, connectivity and genomic span. The detection of cell-type-specific SEs with high centrality within SE communities suggests that key SEs may anchor distinct regulatory sub-networks, possibly coordinating lineage-specific transcriptional states. Additional work is required to functionally characterize the structural and functional roles of SEs with different hierarchical roles in the contact networks. Larger GAM datasets are also required to explore the contributions of different SEs to interchromosomal networks.

### Conclusions and future perspectives

In summary, our findings reveal that SEs form extensive megabase-long range complex chromatin interactions strongly linked with cell-type specific transcriptional programs. The introduction of centrality and community analyses allow us to prioritize SEs according to the hierarchy of their structural and functional roles. To further address the stability and correlation between interaction topologies and the roles of SEs with different hierarchical roles, it will be important to integrate the network topologies with single-cell transcriptomics or live-cell imaging of SE contacts (Cho et al. 2018; Sabari et al. 2018) to further clarify the temporal behaviour, stochasticity, and regulatory impact of SE-interaction networks. Additionally, while the analyses of TF motif enrichment point to putative TF regulators, future work mapping chromatin occupancy of TFs (e.g., ChIP-seq) or CRISPR interference will directly assess their binding and functional relevance for SE formation and SE networks. Another future perspective will be to investigate how disease-associated variants, particularly non-coding variants enriched in SEs, may be involved in complex SE interaction networks, and in that manner expand the discovery of candidate disease genes and pathways. Given the enrichment of central SEs in regulatory hubs, their perturbation may be at the core of network integrity and transcriptional stability, especially in highly specialized cells with fewer SEs that establish more specific networks.

The increased structural complexity and genomic span of SE networks in differentiated neurons suggest a key role for SEs in orchestrating stable, long-range gene regulation in non-dividing terminally differentiated cells. This work provides a blueprint for future investigations into the spatial logic of transcriptional control mediated by 3D genome topology, and prioritizes a set of candidate SEs that may serve as key structural regulators.

## Methods

### Mouse embryonic stem cell culture

Mouse ESCs (46C clone, derived from E14tg2a and expressing GFP under Sox1 promoter; Ying et al., 2003) were a gift from Prof. Domingos Henrique (Instituto de Medicina Molecular, Faculdade Medicina Lisboa, Lisbon, Portugal). The cells were grown as previously described (Ferrai et al. 2017), at 37°C, 5% (v/v) CO_2_ on 0.1% (v/v) gelatine-coated (Sigma, Cat#G1393-100) Nunc^TM^ flasks in Gibco Glasgow’s Minimum Essential Medium (GMEM; Invitrogen, Cat#21710025) supplemented with 10% (v/v) Gibco Foetal Bovine Serum (FBS; Invitrogen, Cat#10270-106, batch number 41F8126K), 0.1 mM β-mercaptoethanol (Invitrogen, Cat#31350-010), 2 mM L-glutamine (Invitrogen, Cat#25030-024), 1 mM sodium pyruvate (Invitrogen, Cat#11360039), 1% penicillin-streptomycin (Invitrogen, Cat#15140122), 1% MEM non-essential amino acids (Invitrogen, Cat#11140035) and 2,000 units/ml Leukemia inhibitory factor (LIF; Millipore, Cat#ESG1107). The medium was changed every day and the cells were split every other day. The mouse ESCs were routinely tested for Mycoplasma contamination using the LookOut® Mycoplasma detection kit (AppliChem, A37440020) according to the manufacturer’s instructions.

Prior to harvesting for RNA extraction, chromatin extraction for ChIP-Seq experiments or to nuclei extraction for ATAC-Seq, the mouse ESCs were grown for 48 hours in serum-free ESGRO Complete Clonal Grade medium (Merck, Cat#SF001-B), supplemented with 1,000 units/ml LIF, on 0.1% (v/v) gelatine-coated Nunc^TM^ dishes (Thermo Fisher, Cat#150326). The medium was changed every 24 hours.

### Dopaminergic neuron differentiation

Mouse ESCs (clone 46C) were differentiated into post-mitotic terminally differentiated dopaminergic neurons as previously described (Jaeger et al. 2011; Ferrai et al. 2017). First, mouse ESCs were differentiated into mouse epiblast-derived stem cells (EpiSCs) by growing the cells for 4 weeks in N2B27 basal medium [1:1 DMEM/F12 (Invitrogen, Cat#21331-020) composed of: Neurobasal medium (Invitrogen, Cat#21103-049), 0.5× N2 (Invitrogen, Cat#17502-048), 0.5x B27 (Invitrogen, Cat#12587-010), 0.05 M β-mercaptoethanol (Invitrogen, Cat#31350-010), 2 mM L-glutamine (Invitrogen, Cat#25030-024)], supplemented with 20 ng/ml Activin (R&D, Cat#338-AC-050) and 12 ng/ml FGF2 (Fibroblast Growth Factor 2; Peprotech, Cat#100-18B). The EpiSCs were grown on 6-well Nunc^TM^ plates coated with FBS (coating for 30 min at 37°C) in a 37°C, 5% (v/v) CO_2_ incubator, and the culture medium was changed every day. The EpiSCs were split every other day by washing the cells 3 times with magnesium– and calcium-free PBS (Phosphate-Buffered Saline), incubating the cells in PBS for 3 min at room temperature and gently scraping the cells from the plate before transferring them to a new FBS-coated plate. EpiSC batches were tested for *Mycoplasma* contamination as previously described for mouse ESCs.

EpiSCs were differentiated into day-30 midbrain DNs using a protocol previously described (Jaeger et al. 2011; Ferrai et al. 2017). One day prior to starting the differentiation, the EpiSCs were grown on 6-well Nunc^TM^ plates coated with 15 μg/ml human plasma fibronectin (Millipore, Cat#FC010) in N2B27 basal medium supplemented with Activin and FGF2, to reach a confluency of 70-80% after 24 h. To start the differentiation process (day 0), the EpiSCs were rinsed twice with magnesium– and calcium-free PBS, N2B27 basal medium supplemented with 1 μM PD 0325901 (Axon, Cat#1408) for 2 days, and the medium was changed every day. On day 2, the cells were washed with magnesium– and calcium-free PBS, scraped, replated on 10 cm Nunc^TM^ plates coated with 15 μg/ml human plasma fibronectin, and cultured in N2B27 basal medium. The cells were kept in N2B27 basal medium for 3 days and half of the medium was replaced every day with freshly prepared medium. After 72 h, the medium was replaced with N2N27 basal medium supplemented with 100 ng/ml FGF8 (Peprotech, Cat#100-25-25) and 200 ng/ml SHH (R&D, Cat#464-sh-025). The cells were cultured for 96 h in N2B27 basal supplemented with FGF8 and SHH medium, and the medium was replaced daily by removing half of the medium and adding half of the volume of freshly prepared medium. After 96 h, the cells were washed once with magnesium– and calcium-free PBS and cultured in N2B27 basal medium supplemented with 10 ng/ml BDNF (R&D, Cat#450-02-10), 10 ng/ml GDNF (Glial Cell-Derived Neurotrophic Factor; R&D, Cat#450-10-10) and 200 μM L-ascorbic acid (Sigma, Cat#A4544) until day 30. Half of the medium was replaced daily with freshly prepared one. To test the quality of the differentiation, cells were collected at different stages of differentiation [mouse ESCs, EpiSCs, day 5 (neuronal progenitor cells, NPCs), day 16 (immature dopaminergic neurons) and day-30 (mature DNs)]. To test the efficiency of the differentiation by whole-cell immunofluorescence, the cells were differentiated in parallel on 22 mm glass coverslips (Thermo Fisher, Cat#150067).

### Cryoblock preparation from cultured dopaminergic neurons

*In vitro* cultured day-30 DNs were fixed and processed into cryoblocks as previously described (Beagrie et al. 2017), using the Tokuyasu method (Tokuyasu 1973). This approach preserves the cellular and nuclear architecture and maximizes the retention of nuclear proteins such as active RNA polymerases (Pombo et al. 1999; Guillot et al. 2004; Branco et al. 2008). The DNs were fixed in 4% and 8% paraformaldehyde (PFA; electron microscopy grade, methanol-free; Alfa Aesar, Cat#43368) in 250 mM HEPES-NaOH (pH 7.6) for 10 min and 2 hours, respectively. The fixed cells were detached from the plates by gentle scraping and collected by centrifugation. As a cryoprotectant that prevents ice crystal damage, the cell pellets were embedded in 2.1 M sucrose (w/v; Sigma-Aldrich, Cat#S9378) in PBS (Sigma-Aldrich, Cat#P44417), assembled on copper stubs and frozen in liquid nitrogen on copper stubs. The cryopreserved samples are stored immersed in liquid nitrogen indefinitely for long-term use.

### Ultrathin cryosectioning

Ultrathin cryosectioning was performed as previously described (Winick-Ng et al., 2021) with a glass knife using an Ultramicrotome (Leica Microsystems, EM UC7) at –110°C. Cryosections of approx. 220 nm thickness were captured in drops of 2.1 M (w/v) sucrose in PBS suspended in a copper wire loop. The captured cryosections were transferred onto 10 mm glass coverslips (thickness nr. 1.5; Marienfeld, Cat#0111500) for confocal imaging, or onto metal framed 4.0 μm polyethylene naphthalate (PEN; Leica Microsystems, Cat#11600289) membranes for laser microdissection.

### Ultrathin cryosectioning and immunofluorescence on PEN membranes

Ultrathin cryosectioning for *immuno*GAM was performed from DNs cryoblocks and the samples were collected on 4.0 μm polyethylene naphthalate (PEN) membranes as described above. For detection of DNs and their nuclei, ultrathin cryosections were co-labelled with sheep anti-Tyrosine Hydroxylase (TH; sheep anti-TH1, Pel Freez, P60101-0) antibodies and mouse anti-pan-histone antibodies H11-4 (mouse anti-pan-histone H11-4, EMD Millipore, MAB3422). After incubation with secondary antibodies, the cryosections were washed twice with PBS, rinsed thrice with ultraclean water (Sigma-Aldrich, Cat#W0502) to remove potential salt crystals, and left to air dry at room temperature for 5 min. To minimize the contamination of the GAM samples, all solutions used for immunofluorescence prior to laser capture microdissection were prepared in ultraclean water and filtered through a 0.21 μm syringe filter (Sartorius, Cat#16555), with the exception of antibody dilutions or stocks.

### Laser capture microdissection

Ultrathin cryosections of DNs were collected on PEN membranes and immunofluorescently labelled to detect neurons and their nuclei. Nuclear profiles (NPs) were microdissected using the Leica LMD7 laser microdissection microscope (Leica Microsystems, LMD7000) using a 63x dry objective. Three NPs per sample were collected into an adhesive PCR cap (Zeiss, Cat#415190-9161-000) and one PCR cap for each 48 PCR cap stripes was left as a negative control (i.e. containing 0 NP). After laser microdissection, samples in PCR caps were stored at –20°C typically overnight or up to a few weeks before proceeding with the next steps.

### Whole-genome amplification

Nuclear profiles were lysed in the PCR adhesive caps for 24 hours at 60°C in 1.2x lysis buffer [30 mM Tris-HCl pH 8.0, 2 mM Ethylenediaminetetraacetic acid (EDTA) pH 8.0, 800 mM guanidium-HCl, 5% Tween 20 (v/v), 0.5% Triton X-100 (v/v)] supplemented with 2x concentration of Qiagen protease (Qiagen, Cat#19155). The protease was inactivated at 75°C for 30 min and the extracted DNA was amplified using random primers with an adaptor sequence. The extracted DNA was pre-amplified using a 2x DeepVent mix (2x Thermo polymerase buffer (10x), 400 μm deoxyribonucleotide triphosphate (dNTPs), 4 mM MgSO_4_ in ultraclean water), 0.5 μM GAT-7N primers (5′-GTG AGT GAT GGT TGA GGT AGT GTG GAG NNN NNN N) and 2 units/μl DeepVent (exo-) DNA polymerase (New England Biolabs, Cat#M0259L) in a thermocycler. To increase the amount of the product a second round of amplification was performed using the primers that annealed to the general adaptor sequence. The amplification step was done using the 2x DeepVent mix supplemented with 10 mM dNTPs, 100 μM GAT-COM primers (5′-GTG AGT GAT GGT TGA GGT AGT GTG GAG) and 2 units/μl DeepVent (exo-) DNA polymerase in a thermocycler (95°C for 3 min, 26 cycles of 95°C for 20 s, 58°C for 30 s, 72°C for 3 min, and one cycle of 72°C for 3 min).

### Library preparation and high-throughput sequencing

After WGA, the samples were purified using solid-phase reversible immobilization beads (1.7 ratio of beads per sample volume) and the concentration of each sample was measured using the Quant-iT^TM^ Pico Green dsDNA assay kit (Invitrogen, Cat#P7589) according to the manufacturer’s instructions. Sequencing libraries were prepared using a Tn5-based in-house protocol adapted from (Picelli et al. 2014). To produce sequencing libraries, Tn5 was first purified and loaded. Briefly, the hyperactive Tn5 protein that contains the E54K and L372P mutations (Adgene, plasmid #60240) was expressed and purified as previously described (Picelli et al., 2014), and stored in 50 mM HEPES-KOH (pH 7.2), 100 mM NaCl, 0.1 mM EDTA, 1 mM dithiothreitol (DTT) and 55% glycerol (v/v). Tn5 expression and purification were performed by the Protein Production and Characterization Platform at the Max Delbrück Center. Tn5 loading was done by mixing 0.36 vol of Tn5 (A280 = 3.0) with a 0.125 vol of a 100 μM equimolar mixture of preannealed Mosaic End double-stranded (MEDS) oligonucleotides [Tn5MEDS-A (5′-TCGTCGGCAGCGTCAG ATGTGTATAAGAGACAG-3′; Illumina, Cat#FC-121-1030) and Tn5MEDS-B (5′-GTCTCGTGGGCTCGGAGATGTGTATAAGAG ACAG-3′), Tn5Merev (5′-[phos]CTGTCTCTTAT ACACATCT-3′)]) in Tn5 dilution buffer (50 mM HEPES pH 7.2, 100 mM NaCl, 0.1 mM EDTA, 1 mM DTT, 0.1% Triton X-100, 50% Glycerol), and incubating the mixture under constant shaking (350 rpm) in a thermomixer at 23°C for 60 min. The primers were pre-annealed prior to loading in annealing buffer (50 mM NaCl, 40 mM Tris-HCl pH 8.0). Tagmentation of GAM samples was done by incubating 1 ng of purified DNA per sample with 5x tagmentation buffer [50 mM TAPS (pH 8.5), 8% PEG, 5 mM MgCl_2_], 10 ng/ul loaded Tn5 in a thermocycler at 55°C for 7 min. The Tn5 inactivation was done by heating the samples at 70°C for 5 min. Next, 1.25 μM Nextera indexes and 0.1X KAPA GC buffer, 0.1 mM dNTPs, 0.1 units/μl KAPA HiFi DNA polymerase (Roche; Cat#KK2102) were added to each tagmented DNA sample. The GAM libraries were prepared by incubating the samples in a thermocycler as follows: 72°C for 3 min, 95°C for 10 s, 12 cycles of 95°C for 10 s, 55°C for 30 s, 72°C for 30 s, and one cycle of 72°C for 5 min.

After library preparation, the DNA was purified using SPRI beads (1.7 ratio of beads per sample volume) and the concentration of each sample was measured using the Quant-iT Pico Green dsDNA assay kit. Equal amounts of DNA from each sample were pooled together (up to 192 samples), and the final pool was purified two more times using SPRI beads (1.7 ratio of beads per sample volume). The final pool of libraries was analysed on an Agilent 2100 Bioanalyzer to confirm the removal of primer dimers and to estimate the average size and DNA fragment size distribution. The libraries were sequenced using the single-end NextSeq 500/550 High Output v2 kit (75 cycles; Illumina, Cat#20024906) on an Illumina NextSeq 500 machine following the manufacturer’s instructions.

### RNA extraction

Total RNA was isolated using TRIzol™ (Invitrogen, Cat#15596026) extraction according to the manufacturer’s instructions. Briefly, TRIzol samples were incubated in chloroform (1:0.2 sample to chloroform ratio) for 3 min at room temperature. After centrifugation (16,260 xg, 15 min, 4°C), the upper aqueous phase was transferred to a new tube and the RNA was precipitated from the aqueous solution using High Performance Liquid Chromatography (HPLC)-grade isopropanol. After centrifugation (16,260 xg, 10 min, 4°C), the RNA pellet was washed with 75% ethanol and eluted in Rnase-free water. The DNA was removed by TURBO™ Dnase (Thermo Fisher, Cat#AM2238) according to the manufacturer’s instructions, and the RNA was stored at –80°C until further processing.

### cDNA synthesis and single gene RNA expression by RT-qPCR

To test and confirm optimal DNA differentiation, samples were collected at key stages throughout the process of differentiation, explained above, and RNA was isolated to test the expression of differentiation markers by quantitative PCR (qPCR) as previously described (Ferrai et al., 2017). Briefly, RNA was reverse transcribed into complementary DNA (cDNA) using random primers and SuperScript™ II Reverse Transcriptase (Invitrogen, Cat#18064-071) according to the manufacturer’s instructions. The cDNA was diluted 1:100 before running qPCR using the 2x SensiMix SYBR No-ROX (Thermo Fisher, Cat#AB1159A) according to the manufacturer’s instructions, and primers that span exon-exon junctions. All amplification products were normalised to the beta actin gene (*Actb*) housekeeping gene.

### Total RNA library preparation and sequencing

Total RNA samples were isolated from ESCs and DNs, and treated with TURBO™ DNase as described above. The quality of the RNA was assessed by running the Agilent Bioanalyzer 6000 RNA Nano assay (Agilent, Cat#5067-1511) according to the manufacturer’s instructions, and only high-quality samples (RIN above 7.30) were further processed. Total RNA libraries were prepared using 1 μg of DNase-treated total RNA and the TruSeq Stranded Total RNA Sample preparation kit (Illumina, Cat#RS-122-2201) according to the manufacturer’s instructions. The quality of the total RNA libraries was assessed before sequencing using the Bioanalyzer High Sensitivity DNA assay (Agilent, Cat#5067-4626). Total RNA-Seq libraries were sequenced using the paired-end NextSeq 500/550 High Output v2 kit (75 cycles; Illumina, Cat#20024906) on an Illumina NextSeq 500 machine following the manufacturer’s instructions.

### Chromatin preparation

Chromatin preparation from DNs was performed as previously described with few modifications (Stock et al. 2007; Brookes et al. 2012; Ferrai et al. 2017). Briefly, cells were fixed using 1% formaldehyde (Sigma-Aldrich, Cat#F8775) for 10 min at 37°C, and the reaction was stopped by adding glycine to a final concentration of 0.125 M and incubating the samples for 5 min at room temperature. The fixed cells were washed three times with ice-cold PBS and lysed in a swelling buffer [25 mM HEPES (pH 7.9), 1.5 mM MgCl_2_, 10 mM KCl, 0.1% NP-40] at 4°C for 10 min. Cells in swelling buffer were scraped from the dish, transferred to a tube, before Dounce homogenization (60 times, ‘tight’ pestle) and purification of nuclei through centrifugation (5,000 rpm, 20 min, 4°C). After extraction, the nuclei were resuspended in sonication buffer [50 mM HEPES (pH 7.9), 140 mM NaCl, 1 mM EDTA, 1% Triton X-100, 0.1% Na-deoxycholate, 0.1% SDS] and sonicated using a Bioruptor Plus® (Diagenode, Cat#B01020001) at 4°C, with the following conditions: high frequency, 60 cycles of 30 s on / 45 s off. The swelling and sonication buffers were supplemented with 5 mM NaF, 2 mM Na_3_VO_4_, 1 mM PMSF, and the cOmplete™ Mini EDTA-free Protease inhibitor cocktail (Roche, Cat#11836170001). After removal of insoluble material by two rounds of centrifugation (4°C, 14,000 rpm, 15 min), chromatin samples were stored at –80°C. Chromatin preparation for mouse ESCs was done as described for DNs, except for the sonication cycles which were as follows: high frequency, 45 cycles of 30 s on / 45 s off. Chromatin concentrations were estimated as previously described (Stock et al., 2007) by measuring absorbance (280 nm) of alkaline-lysed crosslinked chromatin and converting it into arbitrary mass using an estimate of 50 mg/ml for 1 absorbance unit.

### Chromatin immunoprecipitation and sequencing (ChIP-Seq)

ChIP-Seq was done to determine chromatin occupancy profiles for H3K27ac and H3K4me1 in mouse ESCs and DNs. ChIP-Seq experiments were performed as previously described (Stock et al. 2007; Brookes et al. 2012; Ferrai et al. 2017). Briefly, fixed chromatin (350 μg) was immunoprecipitated overnight at 4°C with rabbit anti-histone H3 (acetyl K27) monoclonal antibody (H3K27ac; clone EP16602, Abcam, Cat#Ab177178) or rabbit anti-histone H3 (mono methyl K4) polyclonal antibody (H3K4me1; Abcam, Cat#Ab8895), and 50 μl ChIP-IT^®^ protein G magnetic beads (Active Motif, Cat#53014) in sonication buffer on a rotating wheel. After immunoprecipitation using Immunoglobulin G (IgG) antibodies, the blocked beads were washed once with sonication buffer, once with sonication buffer supplemented with 500 nM NaCl, once with 20 mM Tris (pH 8.0), 1 mM EDTA, 250 mM LiCl, 0.5% NP-40, 0.5% Na-deoxycholate, and twice with TE buffer [1 mM EDTA 10 mM Tris HCl (pH 8.0)]. The immune complexes were eluted from the beads in 50 mM Tris-HCl (pH 8.0), 1 mM EDTA, 1% SDS. Reverse crosslinking was done for 16 h at 65°C in 160 mM NaCl and 20 μg/ml RNAase A (Sigma, Cat#R4642). After increasing the concentration of EDTA to 5 mM the samples were incubated for 2 h at 45°C with 200 μg/ml proteinase K (Roche, Cat#3115836001). DNA was extracted by phenol-chloroform and precipitated by ethanol. The final DNA concentration was measured by Quant-iT PicoGreen dsDNA Assay Kit and Qubit™ 4 (Thermo Fisher, Cat#Q33238) fluorimetry according to the manufacturer’s instructions.

ChIP-Seq libraries were prepared using the TruSeq® ChIP Sample Preparation kit (Illumina, Cat# IP-202-9001DOC) according to the manufacturer’s instructions, except the PCR amplification step was done prior to size selection on an agarose gel (250-600 bp fragments including adaptors). DNA purifications were done using the MinElute PCR purification kit (Qiagen, Cap#28008) and DNA extraction from the agarose gels was performed using the QIAquick Gel Extraction kit (Qiagen, Cat#28704). Libraries were quantified by Qubit and by qPCR using the KAPA Library Quantification Universal kit (Roche, Cat#KK4824). After assessing library sizes by Bioanalyzer with the High Sensitivity DNA analysis kit (Agilent, Cat#5067-4626), the samples were sequenced with the single-end NextSeq 500/550 High Output v2 kit (75 cycles; Illumina, Cat#20024906) on an Illumina NextSeq 500 machine following the manufacturer’s instructions.

### Assay for transposase-accessible chromatin sequencing (ATAC-Seq)

ATAC-Seq for mouse ESCs and DNs was performed according to the Omni-ATAC protocol (Corces et al. 2017), with minor modifications. Briefly, after cell lysis and extraction of the nuclei, the DNA was tagmented in the intact nuclei. Due to the high content of debris in DN samples, manual counting was not as precise as for ESC samples. To overcome this problem, the tagmentation reaction was prepared for three different nuclei dilutions: 1X (50,000 nuclei), 2X and 3X. After running the pre-amplification qPCR, the dilution with the highest TN5-reaction quality was selected. For tagmentation, the nuclei were resuspended in the TAPS-DMF buffer [50 mM TAPS-NaOH (pH 8.5), 25 mM MgCl_2_, 50% DMF], 0.1% Tween-20, 0.1% digitonin, in 0.25x PBS. A total of 3 μl of Tn5 mix (5.6 μg Tn5 and 0.143 volume of 100 μM adapter mix) was added to the transposition reaction mix. The tagmented DNA was purified, and the optimal number of cycles for amplification and the sample quality were assessed by qPCR. Libraries were generated according to the Omni-ATAC protocol using the Illumina index 1 (i7) and index 2 (i5) adapters. The ATAC-Seq samples were sequenced with the paired-end NextSeq 500/550 High Output v2.5 kit (150 cycles; Illumina, Cat#20024907) on an Illumina NextSeq500 machine following the manufacturer’s instructions.

### RNAPII-S5p RNA ChIP-Seq

To capture chromatin-bound RNA in co-association with RNAPII-S5p, immunoprecipitation was performed using a mouse anti-RNAPII-S5p antibody (clone CTD4H8, BioLegend, Cat#904001), which detects RNA polymerase II throughout coding regions of active genes (Brookes et al. 2012). Briefly, the protein-G-magnetic beads were incubated with rabbit anti-mouse (IgG+IgM) bridging antibodies (Jackson Immunoresearch, Cat#315-005-048), 10 μg antibodies per 50 μl beads, for 1 h at 4°C, before washing the sonication buffer. The remaining steps before reverse crosslinking were carried out as described above for ChIP-Seq. To prevent RNA degradation, 50 units/ml RNaseOUT™ recombinant ribonuclease inhibitors were added to the immunoprecipitation reagents.

Reverse cross-linking was done to extract chromatin bound RNAs, by adding NaCl to a final concentration of 200 mM and incubating the samples for 4 h at 65°C. After increasing the EDTA to a final concentration of 5 mM, samples were incubated in 200 μg/ml proteinase K for 1 h at 40°C. The RNA was recovered by acid phenol-chloroform extraction and isopropanol precipitation. After TURBO™ DNase treatment and quantification, the RNA was reverse transcribed into cDNA and the quality of the immunoprecipitated RNA was first assessed by qPCR. To retain information about the direction of the transcription, directional RNA-ChIP-Seq libraries were prepared from the immunoprecipitated RNA using the Illumina small RNA sample preparation kit with the v1.5 sRNA 3’ adaptor (Illumina, Cat#FC-930-1501), according to the manufacturer’s instructions. Libraries were quantified using Qubit and the fragment sizes were assessed with the Bioanalyzer. The samples were sequenced on an Illumina HiSeq machine according to manufacturer’s instructions.

### Mapping immunoGAM DNs data and calling positive windows

Mapping and calling positive windows for the DN immunoGAM data was done as previously described (Winick-Ng et al. 2021). Raw sequencing reads for the immunoGAM DNs data were mapped to the mouse genome assembly GRCm38 (mm10, December 2011) using Bowtie2 (v.2.3.4.3) with the default settings (Langmead and Salzberg 2012). Reads with mapping quality below 20, non-uniquely mapped reads and PCR duplicates were excluded. Positive genomic windows present in the ultrathin NPs were identified for individual immunoGAM libraries. For each resolution used, the genome was split into equal-sized windows and the number of nucleotides sequenced per bin was calculated using BedTools v2.30.0. The percentage of orphan windows (i.e. positive windows flanked by two adjacent negative windows) was calculated for every percentile of the nucleotide coverage distribution. The optimal threshold for identifying windows in each immunoGAM sample was calculated as the number of nucleotides corresponding to the percentile with the lowest percentage of orphan windows. A window was called positive if the number of nucleotides sequenced in each bin was greater than the optimal threshold.

### immunoGAM quality control metrics

Quality controls for each dataset were done by determining the percentage of orphan windows in each sample, the number of uniquely mapped reads to the mouse genome and the correlations from cross-well contamination for every sample (**Supplemental Figure S1D**). Samples that passed the quality control criteria had < 60% orphan windows, > 50,000 uniquely mapped reads to the mouse genome and a cross-well contamination score per collection plate < 0.4 Jaccard index.

### Processing of published mouse ESCs GAM data

The published mouse ESCs GAM datasets used were remapped to the mouse genome assembly GRCm38 (mm10, December 2011) and processed to call positive windows using a negative binomial approach as previously described for GAM data collected with a previous genomics pipeline based on scWGA kit from Sigma (Beagrie et al. 2017; Beagrie et al. 2023). Published GAM datasets were available from two biological replicates and were collected in two modes, from single nuclear profiles (1NP) or using multiplex-GAM that combines 3NPs in one GAM reaction. To obtain GAM dataset in mouse ESCs equivalent to the 3NP multiplex GAM dataset collected here from DNs, the published 1NP GAM datasets that passed quality control were converted, *in silico*, into 3NP GAM samples by random pooling three 1NP GAM datasets into in-silico 3NP GAM samples, as previously (Beagrie et al. 2023).

### Curating the immunoGAM datasets

After selecting the samples that pass the quality control criteria, the whole GAM dataset was further curated to remove windows that are over-or under-detected, for example due to low read mapping, for example due to repetitive and redundant regions. The over– and under-detected windows were removed from the GAM datasets according to (Irastorza-Azcarate et al. 2025). Briefly, for each resolution and dataset a signal smoothing algorithm (i.e. smoothing variable of 11 windows) was applied to the window detection frequency (WDF) and, for each genomic window, the fold change between the raw and smoothed WDF was computed. Windows with a fold change between raw and smoothed higher than 5 were not considered in downstream analyses. Finally, mappability data were generated for each resolution using the GEM-mappability software (https://evodify.com/gem-mappability/) over the indexed mouse genome assembly GRCm38 (mm10, December 2011) and 75-kmer size. The mean mappability score was then computed for each genomic window at a given resolution using the bigWigAverageOverBed utility from UCSC tools (https://genome.ucsc.edu/ goldenPath/help/bigWig.html). Windows that had a mean mappability score > 0.2 were discarded.

### Visualizing pairwise GAM contact matrices

For visualization of GAM data, contact matrices for mouse ESCs and DNs were calculated using pointwise mutual information (PMI) for all pairs of windows genome-wide, as previously described (Winick-Ng et al. 2021). The PMI for given genomic regions *I* and *j* describes the difference between the probability (p) of their joint distribution (p(*I,j*), i.e. both windows being found in the same NP) and their individual distributions (p(*i*) and p(*j*)) across all NPs. PMI was calculated using the following formula: PMI = log(p(*I,j*) / p(*i*) p(*j*)). To produce a normalised PMI (NPMI), the PMI was restricted between –1 and 1 using the following formula: NPMI = PMI / –log(p(*I,j*)). For visualization of contact matrices, the scale bars were adjusted in each genomic region displayed to a range between 0 and the 99^th^ percentile of the NPMI values for each GAM dataset.

### Insulation score and topological associating domain boundary calling

TADs were defined in ESCs and DNs GAM data using the insulation score method as previously described (Crane et al. 2015; Winick-Ng et al. 2021; Beagrie et al. 2023). Briefly, insulation scores for each dataset were calculated on NPMI GAM contact matrices at 50 kb resolution using insulation square sizes ranging from 100 to 1000 kb. TAD boundaries were called using a 500 kb insulation square size and defined as the local minima of the insulation score. Boundaries that touched or overlapped by at least one nucleotide were merged.

### Extracting significant pairwise and triplet contacts with SLICE

The SLICE model was applied, as previously described for multiplex-GAM data (Beagrie et al. 2017; Beagrie et al. 2023). Briefly, SLICE was used to extract intra-chromosomal pairwise (Pi2) and three-way (triplet) interaction probabilities (Pi3) from curated GAM co-segregation data at 5 kb and 150 kb, respectively. The following parameters were used as input: mouse genome assembly GRCm38 (mm10, December 2011), mouse diploid genome length (G) of 5.46 Gbp, nuclear slice thickness (h) of 0.22 μm; nuclear radius of mouse ESCs of 4.60 μm (Beagrie et al. 2017) and nuclear radius of DNs of 3.25 μm, obtained as previously described (Beagrie et al. 2017).

### Processing and analysis of total RNA-Sequencing data

Four biological replicates of ESCs and four of DNs were used for the differential expression analysis. For both ESCs and DNs, two of the libraries were sequenced ultra-deep. To obtain sequencing datasets of similar read depth for all biological replicates, the samples sequenced ultra-deep (sample identifiers: esc_A5, dopa30_A1, dopa30_A2) were subsampled to the same number of reads as the next largest sequencing file (60,301,920 for ESCs and 93,104,024 for DNs) with seqtk v1.3-r10 (https://github.com/lh3/seqtk) using 123 as a random seed. The RNA-Seq data was mapped to the Ensembl GRCm38 v.102 list of annotated transcripts using STAR v2.6.0c (Dobin et al. 2013) and the annotated genes were quantified with RSEM v1.3.0 (Li and Dewey 2011). Bigwig tracks for visualization in the genome browser were produced with the deepTools v3.1.3 command bamCoverage (Ramírez et al. 2016) with base per million (BPM) normalization, after sorting and indexing of the bam files with samtools v1.12 (Danecek et al. 2021). Differential expression analysis was performed using the DESeq2 v1.2.0 (Love et al. 2014) for genes detected above 1 transcript per million (TPM) in at least one sample. Genes that had a log2(Fold Change) > |2|, an adjusted p-value < 0.05 and a delta TPM (|ESCs_meanTPM – DNs_meanTPM|) > 1 were classified as differentially expressed genes.

### Processing of the ChIP-Seq data

Processing of the ChIP-Seq libraries for H3K27ac and H3K4me1 was performed as previously described (Ferrai et al. 2017). Briefly, sequenced reads were aligned to the mouse genome assembly GRCm38 (mm10, December 2011) using Bowtie2 v2.0.5 (Langmead and Salzberg 2012) with default parameters. Identical reads that aligned to the same genomic location (i.e. duplicated reads) occurring more often than the 95^th^ percentile of the frequency distribution of each dataset were removed using a custom pipeline.

### ChIP-Seq data visualization

Bigwig tracks were generated using Bedtools v2.30.0 genomeCoverageBed program with the standard parameters. The bedgraph files were then converted to bigwig files using UCSC tools wigToBigWig utility (Kent et al., 2010). The UCSC Genome Browser (http://genome.ucsc.edu) was used to visualise ChIP-Seq data, using default settings, with the following exceptions: “Window function” = “mean”, “Smoothing Window” = “2 pixels”, and inclusion of zero on the y-axis.

### Peak calling in ChIP-Seq data

For H3K4me1 and H3K27ac, MACS2-broad performed best in mouse ESCs, whilst HOMER performed best in DNs (Heinz et al. 2010). HOMER and MACS2-broad algorithms were run using the default parameters, using ChIP-Seq data produced using anti-digoxigen from ESCs and day-16 *in vitro* differentiated DNs (GSE94364) (Ferrai et al. 2017), as control background datasets for ESCs and DNs, respectively.

### Mapping and processing of the ATAC-Seq data

The ATAC-Seq Fastq files were mapped to the mouse genome assembly GRCm38 (mm10, December 2011) using Bowtie2 v2.3.4.3, with the default parameters except: – –local, – –minions 25, –-maxins 2000 – –no-mixed, –-no-discordant –dovetail –soft-clipped-unmapped-tlen. Reads mapping to the mitochondrial genome and low-quality reads (MQ < 30) were discarded. The sequencing files were then sorted and the duplicated reads were removed using Sambamba v0.6.8 (Tarasov et al. 2015) markdup with default parameters except: –-remove-duplicates, –-hash-table-size 525000, –-overflow-list-size 525000. For further analysis, BAM files were filtered out to remove reads mapping to non-canonical chromosomes (i.e. only reads mapping to chr1 to chr19, chrX and chrY were included). Reads per kilobase of transcript, per million mapped reads (RPKM) tracks were generated using the deepTools bamCoverage tool v3.1.3 with ‘binsize’ = 1 (Ramírez et al. 2016).

### Peak calling in ATAC-Seq data

Peaks were called for each ATAC-Seq sample using MACS2 v2.1.1.20160309 (Zhang et al. 2008). Two sets of peaks were generated (i.e. broad and narrow) for each replicate using the corresponding algorithms and the following parameters: –-nomodel –shift – 100 –extsize 200 –bdg –SPMR –keep-dup all –qvalue 0.05.

### Processing of the RNAPII-S5p RNA ChIP-Seq

Sequenced paired-end reads originating from RNAPII-S5p RNA ChIP-Seq libraries were mapped to the indexed mouse genome assembly GRCm38 (mm10, December 2011), using as reference the Ensembl GRCm38 v.102 list of annotated transcripts and the STAR v2.7.8a software (Dobin et al. 2013). Aligned reads were filtered to keep only uniquely mapped reads and in proper pairs, using samtools v1.12 (Danecek et al. 2021). Reads originating from forward and reverse strands were separated using samtools v1.12 view (Danecek et al. 2021). Reads that were first-in-pair and mapped to the reverse strand or second-in-pair and mapped to the forward strand were assigned as originating from the forward strand. Reads that were first-in-pair and mapped to the forward strand and second-in-pair and mapped to the reverse strand were assigned as originating from the reverse strand. To visualize RNAPII-S5p RNA ChIP-Seq data in UCSC, bigwig tracks were first generated using Bedtools v2.30.0 genomeCoverageBed and then converted to bigwig using UCSC tools wigToBigWig utility (Kent et al. 2010).

### Calling of super-enhancers

To call super-enhancers (SEs), robust ATAC-Seq MACS2-broad peaks that did not overlap known promoters (i.e. the 2.5 kb window centered around each TSS) were selected and classified based on their overlap with ChIP-Seq data as described above. Peaks classified as active putative (i.e. overlapped with H3K27ac and H3K4me1 and not H327me3) were used as an input for the ROSE pipeline to call SEs (Hnisz et al. 2013; Lovén et al. 2013; Whyte et al. 2013). First, the list of putative constitutive putative enhancers was converted to a compatible GFF file format. The ROSE_main.py pipeline was run on the GFF file using the H3K27ac and ATAC-Seq BAM files as inputs (-r and –b flags, respectively) and the corresponding digoxigenin ChIP-Seq data as control (-c). To generate the final list of SEs, putative constitutive enhancers within a 12.5 kb window were stitched together, ranked according to their normalised H3K27ac signal and classified as SEs using the tangent of the curve as previously described (Whyte et al. 2013).

### Gene ontology enrichment analysis

Gene ontology (GO) enrichment analysis for different classes of genes was performed with WebGestalt (WEB-based Gene SeT AnaLysis Toolkit) v2019 (http://www.webgestalt.org; Wang et al. 2017) for *Mus musculus*, using Over-Representation Analysis (ORA), default parameters and the backgrounds specified in the text.

### Enrichment analysis of pairwise and triplet SLICE data

Analysis to determine whether prominent pairwise or triplet interactions detected by SLICE were enriched for particular features was performed as previously described, but at higher genomic resolutions (Beagrie et al. 2017; Beagrie et al. 2023). First, the genome was binned into 5 kb bins or 150 kb bins which were then classified based on their overlap with features of interest (e.g. nascent transcription, classes of genes, enhancers, SEs) using BedTools v2.30.0. Bins that overlapped more than one feature were classified as mixed, to avoid ambiguity. The number of observed contacts was calculated by counting the number of times bins classified based on the given features were identified in prominent SLICE pairwise or triplet interaction. As a random control, the list of contacts was permuted 1,000 times by shifting all their genomic positions by a given random distance. If the permuted values were larger or smaller than the chromosome length their position was wrapped around to the end or the beginning, respectively, of the chromosome. This approach preserved the number of contacts per chromosome and their distance distribution. After classifying the bins in the permuted interactions, the number of expected pairs or triplets was counted. P-values were calculated as the proportion of expected values that have a test statistic greater than that of the observed values divided by the number of permutations plus one (Phipson and Smyth 2010).

### Statistical methods and plotting

Statistical analyses were performed using the SciPy v1.9.3 module from Python (Virtanen et al. 2020), and the statistical tests used are mentioned in each figure legend. All plots presented here were produced in Python v3.8.2 using matplotlib v3.6, plotly v5.11.0 or seaborn v0.12.1 modules (Hunter 2007; Waskom 2021).

### TF motif analysis

The motif-analysis tool from Regulatory Genomics Toolbox (Li et al. 2023) was used on SE summits +/-250 bp (6,148 ESC SE summits, 5,536 DN SE summits) with an FDR threshold of < 1×e^−5^. We used annotated HOCOMOCOv12 mouse motifs of A, B, and C quality (Vorontsov et al. 2023) corresponding to 354 total expressed TFs from ESCs or DNs as our motif database. TF motif occurrences for SEs were determined by adding together all that contained SE summit TF presence. TF motif frequency was compared across 754 ESC and 465 DN unique SEs that contained at least 1 TF motif from HOCOMOCO using the two-sided independent samples t-test from SciPy 1.14.1 (Virtanen et al. 2020). Similarly, common SE TF motif distributions were compared (< 0.05 p-value threshold).

For all cell types individually, SEs were separated into 2 groups based on whether they interacted distally and locally (> 1 Mb and ≤ 1 Mb) or only distally (> 1 Mb), and expressed TF motif density p-values were calculated as previously mentioned. DEGs were categorized in the same way, and motif density was recorded for coding regions and compared using the same method.

### Constructing SE-SE interaction networks

We constructed cell-type specific, intrachromosomal SE interaction networks from the top 2% of SLICE triplets that contained an SE at each anchor (12,025 ESC triplets, 2,675 DN triplets). Interacting windows were mapped to intersecting SEs. Edges were mapped such that every node in the network corresponds to one SE, and all SE interactions were preserved. The median of all contributing SLICE *Pi3* values that contained a given pair of SEs was used for the edge weights. Next, we identified central nodes across the ESCs and DNs separately. A centrality ensemble of 8 different measures – Katz, Laplacian, Unweighted/Weighted Degree Centrality, Eigenvector, Harmonic, Betweenness, Closeness – was applied to the SE interaction networks to select the top 2 nodes per chromosome by every measure. All centrality measures were applied from NetworkX 3.4.2 without any modifications (Hagberg et al. 2008), except for weighted degree centrality, which was calculated by the following equation: DC(i) / (n-1), where *DC(i)* is the degree centrality for a weighted graph (Zuo et al. 2012) and (*n-1)* is the number of nodes in the graph, excluding node *i*.

We identified communities of strongly interacting SEs in our cell-type specific intra-chromosomal networks. Python packages igraph 0.11.8 and leidenalg 0.10.2 (Csárdi and Nepusz 2006; Traag et al. 2019) were used in each interaction network with the default RBConfigurationVertexPartition partition method and a resolution of 1.0. Next, by using the igraph 0.11.8 and matplotlib.pyplot 3.8.0 python packages (Csárdi and Nepusz 2006; Hunter 2007), we displayed networks of specific chromosomes for each cell type, highlighting in different levels the nodes selected as central nodes according to our centrality ensemble method. Finally, to identify communities with highly central SEs, communities in each cell type were ranked by strongest central node percentage: (number of central nodes in a community) / (total nodes in a chromosome). SLICE triplets involved with the SE interaction network were plotted in the WashU Epigenome Browser along with a track of the central nodes for each cell type (Li et al. 2022).

### TF pair Networks

To analyse TF interaction activity in the two cell types, edges were extracted from each SE interaction network and all possible pairs of expressed TFs were generated. For every pair, the number of edges were counted when the pair was present in SE-SE interactions. For a given pair {TF_1_, TF_2_}, we define TF pair presence, *P*, in an SE-SE interaction {SE_1_, SE_2_} with the following relation: *P* = (TF_1_ SE_1_ TF_2_ SE_2_) (TF_2_ SE_1_ TF_1_ SE_2_). Cytoscape 3.10.2 (Shannon et al. 2003) was used to display the TF pairs with high interaction enrichment (>40%) as a network, where each node is a TF, and edge weights correspond to the number of contributing SE-SE interaction network edges. The size of every node is scaled based on how many edges contain the TF represented by the node. Node colour indicates Leiden community membership generated akin to SE interaction networks.

### Genomic Distances

To evaluate the genomic distances of the SEs in the constructed network, we started by extracting all possible distances in SE triplets and building every possible triplet in a given chromosome for each cell type and extracting two linear distances – between the first and second SEs and between the second and third SEs in a triplet, ordered according to their genomic coordinates on a chromosome. This served as our control group, as it would indicate all distances that could be recovered if every SE was connected to every other SE. We also evaluated the distances of only those SEs in each cell type that were selected as a central node by our centrality ensemble method. All data was stored/manipulated in data frames using the Python package pandas version 2.2.3 (McKinney 2010).

By using the matplotlib.pyplot 3.8.0 python package (Hunter 2007), we generated density plots where the larger distance in every triplet was located on the y-axis and the shorter distance in every triplet was located on the x-axis, and every triplet was mapped accordingly. Each density plot maps a given group of triplets of a given cell type. This way, we observed the genomic distances of all possible triplets or only central nodes in either DNs or ESCs. In addition to the density plots, we utilized the same data through the same python package to develop box plots evaluating the ranges and the median larger distance of a triplet for each density plot created.

## Data access

All raw and processed sequencing data generated in this study have been submitted to the NCBI Gene Expression Omnibus (GEO; www.ncbi.nlm.nih.gov) under superseries accession number GSE305152. ATAC-Seq data can be found under accession number GSE304720. ChIP-Seq data can be found under accession number GSE304728. GAM and SLICE data can be found under accession number GSE304996. RNA ChIP-Seq can be found under accession number GSE305012. RNA-Seq data can be found under accession number GSE305150. Supplemental Data files are available at Zenodo (https://doi.org/10.5281/zenodo.16780836). Scripts are provided as Supplemental Code on GitHub (github.com/pombolab/Harabula_Speakman_Superenhancer_networks_2025).

## Acknowledgements

A.P. acknowledges support from the Helmholtz Association, the Deutsche Forschungsgemeinschaft (DFG; German Research Foundation) under Germany’s Excellence Strategy [EXC-2049–390688087], and the Medical Research Council [MRC, UK; grant MC_U120061476]. A.P. and I.H. acknowledge support from the DFG [priority program SPP2202]. A.P. and L.Z.R. acknowledge support from the DFG [International Research Training Group IRTG2403]. A.P. and M.N. acknowledge support from the NIH Common Fund 4D Nucleome Program grant 1UM1HG011585-03. L.W. acknowledges support from the iSURF and Stocker Fellowship programs of the School of EECS at Ohio University. I.H. was supported by a Boehringer Ingelheim Fonds PhD fellowship. L.S. was supported by the National Science Foundation Graduate Research Fellowship under Grant No. 2334422 (to L.S.). S.C. was supported by the Fundação para a Ciência e Tecnologia (PD/BD/135453/2017 and COVID/BD/152489/2022). I.I.-A was supported by a Long-Term Fellowship from the Federation of European Biochemical Societies (FEBS).

## Author contributions

Conceptualization: I.H., A.P., L.W., M.N., L.S.; Data curation: I.H., L.S., A.K., I.I.A.; Formal analysis: I.H., L.S., F.M., L.Fio., L.Z.R., A.K., R.B., L.Fer., I.I.-A, D.S.; Funding acquisition: A.P., I.H., M.N., L.W., S.C., I.I.-A.; Investigation: I.H., L.Z.R., A.K., R.B., A.M.F., S.C., C.F.; Methodology: I.H., L.S., F.M., L.Fio., L.Fer., K.J.M., M.N., L.W., A.P.; Project administration: I.H., A.P.; Resources: A.P., M.N., L.W.; Software: I.H., L.S., F.M., L.F., L.Z.R., A.K., L.Fer., I.I.-A. M.N., L.W.; Supervision: A.P., L.W., I.H., M.N.; Validation: I.H., L.S., F.M., L.Fio., M.N., L.W., A.P.; Visualisation: I.H., L.S., L.Fer., A.P.; Writing – original draft: I.H., A.P., L.S., L.Fer.; Writing – review or/and editing: I.H., L.S., L.Z.R., A.K., L.Fer., S.C., D.S., C.F., L.W., A.P.

